# Paired-surface spatial mechanomics links tissue stiffness maps to spatial transcriptomics

**DOI:** 10.64898/2026.08.29.748050

**Authors:** Huan Ting Ong, Yuting Lou, Jake Turley, Ranmadusha M. Hengst, Md Faris H. Ramli, Xingyu Shen, Jennifer Marlena, Jin Zhu, Rong Li, Chii Jou Chan, Jennifer L. Young

## Abstract

Tissue mechanics influence diverse biological processes, yet directly linking stiffness measurements to spatially resolved molecular states in intact tissues remains challenging. Here we developed a paired-surface spatial mechanomics approach to map Young’s modulus by nanoindentation on a fresh tissue surface and co-register the stiffness grid with Visium HD spatial transcriptome data from the immediately adjacent, parallel surface. This was applied to the mouse ovary, which has spatially distinct compartments and undergoes extracellular matrix remodeling with age. Nanoindentation at 50 µm grid spacing enabled millimeter-scale stiffness maps while balancing acquisition time in fresh tissues, generating 2,771 matched measurements across 21 regions of interest. Global and compartment-specific analyses associated stiffer regions with lower matrix-related programs and higher cell-cycle programs, alongside age-dependent inflammatory and metabolic associations. This correlative strategy integrates experimentally measured mechanics with spatial omics in fresh tissues.

## Main text

Tissues are defined not only by their cellular and molecular composition but also by the physical properties of their microenvironment. The extracellular matrix (ECM) provides structural support, transmits forces, and remodels across development, aging and disease, while its mechanical properties and architecture affect cell fate and tissue homeostasis (1,2). Spatial transcriptomics and proteomics now resolve molecular states at cellular scale, yet the physical context in which these states arise is rarely measured. Mechanical and molecular maps are therefore generally obtained from separate samples or regions, preventing direct tests of whether local molecular programs vary with the local mechanical environment.

Available approaches that map tissue mechanics trade spatial resolution against coverage area, acquisition time, and sample processing. Atomic force microscopy (AFM) can resolve mechanics at cellular and matrix fiber scales and has revealed heterogeneous mechanical landscapes in tumors and ovaries (3–5), but high-resolution mapping across millimeter-scale tissues remains limited (4). Large probe nanoindentation increases coverage while averaging mechanics over larger. micron-scale regions (6). Micro-elastography recovers 3D elasticity without contact, but relating these measurements to spatial molecular features is challenging (7). The main obstacles in bridging highly sensitive omics assays with mechanical measurements are acquisition time, sample degradation, and incompatible processing workflows. These approaches therefore provide spatial mechanical maps, but not molecular profiles at matched measurement coordinates.

Several strategies have begun to bridge this gap. STIFMap uses AFM-trained collagen and nuclear image features to predict stromal elasticity and register it with protein biomarkers in breast tumors (8). Force inference has been integrated with spatial transcriptomics to relate epithelial interfacial tension to gene expression (9), while a bone mechanomics platform combines spatial transcriptomics with micro-computed tomography-derived finite element strain (10). Together, these studies establish mechanics as a spatial data layer to molecular analyses and demonstrate the feasibility of linking tissue stiffness with gene or protein expression. However, a workflow that directly co-registers experimentally measured stiffness values of large area mechanical maps in fresh, hydrated tissues with high-resolution spatial transcriptomic profiles remains needed, particularly in heterogeneous, ECM-rich tissues.

The ovary provides a demanding test of such a method because it undergoes extensive, recurrent matrix remodeling. In the ovary, the surrounding stroma remodels as follicles grow, ovulation ruptures and initiates repair at the tissue surface (11,12), and corpora lutea form and regress during each reproductive cycle. Mechanical perturbations regulate follicle activation and oocyte dormancy (13,14), while reproductive aging is accompanied by stromal stiffening, collagen accumulation, altered fibroblast and immune populations, and impaired tissue remodeling (6, 15–19).

This dynamic and spatially heterogeneous tissue architecture makes regional mechanical measurements strongly dependent on where the tissue is sampled. Although ovarian mechanics (3, 4, 6, 7, 19–22) and molecular features of aging (23–31) have each been mapped independently, whether local stiffness corresponds to local transcriptional state across ovarian tissues remains unknown.

Here we establish a correlative spatial mechanomics workflow that co-registers tissue mechanics with the spatial transcriptome and supports both global and compartment-specific analyses (Fig. 1). Fresh vibratome sections of tissues are indented within hours of harvest, while the tissue block immediately beneath and parallel to the indented surface is cryopreserved for spatial transcriptomic capture. Matched vascular landmarks set the orientation and scale of the paired surfaces, while grid reconstruction places the nanoindentation measurements within a common coordinate frame, allowing local stiffness values to be related to gene expression and tissue architecture. Applied to reproductively young and aged mouse ovaries, this approach identifies associations between local stiffness and matrix, metabolic, cell cycle, and immune programs, with different patterns across age groups and tissue compartments. This paired-surface strategy provides a method for correlating experimentally measured mechanics with spatial omics in fresh, sectionable tissues. Ultimately, by bridging physical and molecular data, our correlative spatial mechanomics approach expands the scope of multi-omics analyses and offers a route to understanding tissue function, aging, and disease.

**Figure 1.**
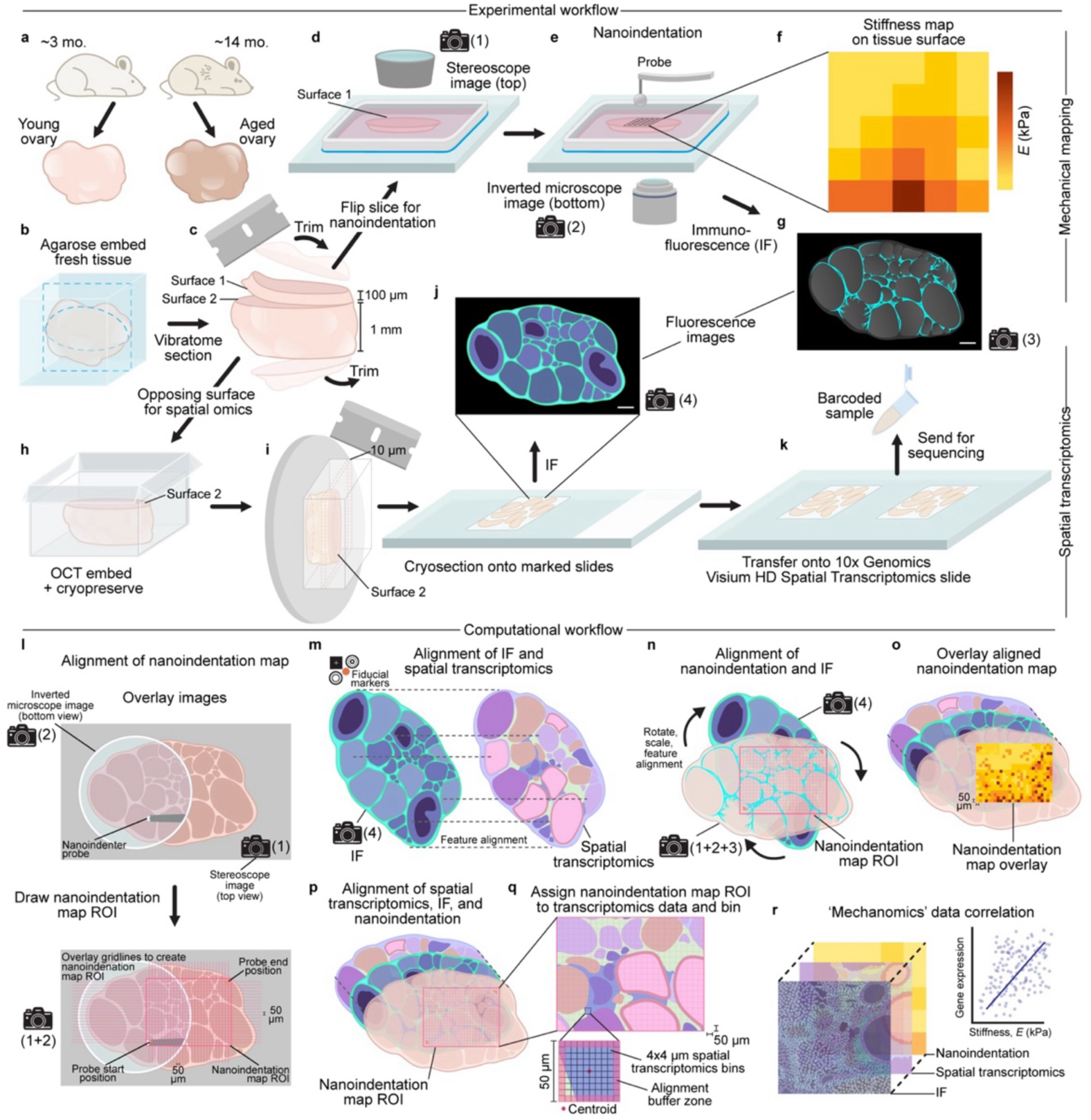
Paired mechanical and transcriptomic measurements across adjacent ovarian tissue surfaces. (**a**) Ovaries were isolated from reproductively young (∼3 months) and aged (∼14 months) CD-1 mice. (**b**) Fresh ovaries were embedded in low-melting-point agarose. (**c**) A 100 µm vibratome slice was generated, producing two opposing surfaces. Surface 1 of the section was used nanoindentation, while surface 2 of the remaining 1 mm tissue block was retained for spatial transcriptomics. (**d**) The vibratome section was flipped, attached to a poly-D-lysine-coated slide under L-15 medium, and imaged from above using a stereomicroscope (image 1). (**e**) Nanoindentation was performed at 50 µm center-to-center grid spacing, with the probe and tissue imaged from below using an inverted microscope (image 2). (**f**) Force-displacement curves yielded stiffness (Young’s modulus, *E*), at each grid point, generating a stiffness map of the indented surface. (**g**) After nanoindentation, the same section was then fixed, immunostained with the same antibody panel used for spatial transcriptomics sections, and imaged by confocal microscopy (image 3). (**h**) The opposing block was embedded in OCT and fresh frozen with surface 2 facing the cutting plane. (**i**) The block was cryosectioned at 10 µm thickness onto marked slides. (**j**) Sections were stained via IF and imaged on a slide scanner (image 4). (**k**) The sections were then transferred onto 10x Genomics Visium HD slides via CytAssist and processed for sequencing. (**l**) To reconstruct the 50 µm-spaced grid on the nanoindented tissue section, images 1 and 2 were overlaid to locate the initial and final probe positions to define the nanoindentation ROI. (**m**) The cryosection IF (image 4) was aligned with the Visium HD spatial transcriptome data using fiducial markers and shared tissue features. (**n**) The post-nanoindentation IF image (image 3), together with images 1 and 2, was rotated, scaled, and aligned with image 4 using corresponding αSMA-positive vascular features, placing the nanoindentation ROI in the coordinate system of the opposing surface. (**o**) The transformed stiffness map was then overlaid on the opposing tissue surface. (**p**) The nanoindentation, IF, and spatial transcriptomic layers were combined in a common coordinate space (**q**) The 4 x 4 µm spatial transcriptome bins were assigned to each reconstructed 50 µm nanoindentation grid point, retaining up to the 140 bins nearest its centroid and excluding the peripheral alignment buffer region. Each grid point contained a median of 89 bins. (**r**) Gene counts were aggregated per grid point and related to the Young’s modulus measured at the corresponding indentation position. Schematics are not to scale.

## Results

### Paired-surface spatial mechanomics links fresh tissue nanoindentation with Visium HD spatial transcriptomics

We developed a paired-surface spatial mechanomics workflow that correlates mechanical measurements acquired from fresh tissue to spatial gene expression from the immediately adjacent, parallel surface. Ovaries were harvested from reproductively young and aged CD-1 mice (Fig. 1a) and embedded in low-melting-point agarose (Fig. 1b). To obtain paired surfaces, a single vibratome cut was made in an isolated ovary, and the two opposing faces were separately assigned to the two modalities (Fig. 1c). The 100 µm vibratome tissue section was attached to a coated slide and nanoindented within a chamber while immersed in medium, with top-view stereomicroscope and bottom-view inverted microscope images recording the entire tissue piece and probe positions, respectively (Fig. 1d,e; Supplementary Methods S1). Nanoindentation yielded a stiffness map of the indented surface (Fig. 1f), after which the section was fixed, stained, and imaged by confocal microscopy using the same immunofluorescence (IF) panel applied to the opposing surface (Fig. 1g). The opposing face of the remaining tissue block was embedded in OCT and cryopreserved (Fig. 1h), cryosectioned at 10 µm thickness (Fig. 1i), stained for IF (Fig. 1j), and transferred onto a 10x Genomics Visium HD spatial transcriptomics slide using CytAssist for subsequent sequencing (Fig. 1k).

To carry out alignment of the paired surfaces, we integrated the four images collected across the two workflows (worked example shown in Extended Data Fig. 1a-c). First, the top-view stereomicroscope image of the nanoindented section and bottom-view inverted microscope images were overlaid to locate the initial and final probe positions, define the nanoindentation region of interest (ROI), and reconstruct the 50 µm spacing grid (Fig. 1l). Second, the IF image from the opposing cryosection was aligned with the spatial transcriptome data using the Visium HD fiducial markers and shared tissue features (Fig. 1m). Third, the post-nanoindentation IF image was aligned with the opposing-surface IF image in SpatialX software by rotation, scaling, and matching α SMA-positive vascular features, placing the nanoindentation ROI in the coordinate system of the opposing surface (Fig. 1n). The transformed stiffness map was then overlaid on the opposing tissue surface (Fig. 1o), and the nanoindentation, IF, and spatial transcriptomic layers were combined within a common coordinate frame (Fig. 1p). Finally, 4 µm spatial transcriptome bins were assigned to the reconstructed nanoindentation grid points (Fig. 1q), enabling stiffness-gene expression analyses (Fig. 1r).

After filtering coordinate and gene expression data, 2,771 stiffness grid points were retained across 21 ROIs from eight ovaries of six mice. Each stiffness grid point carried a median of 89 4-µm spatial transcriptome bins. Bins projecting withing each reconstructed grid cell were retained up to a maximum of the 140 bins nearest its centroid, thereby excluding bins at the edges, where assignment is most sensitive to residual positioning error (Fig. 1q). Assigned bins inherited the stiffness value of the corresponding grid point for spatial visualization, while gene counts were aggregated at the 50 µm grid point, which served as the unit of observation for all subsequent correlations (Fig. 1r).

Correspondence between the two surfaces was established from tissue features rather than from an assumed geometric relationship. αSMA-positive vascular features near the cut plane were resolved on both opposing surfaces, providing landmarks distributed throughout the tissue rather than relying only on the tissue outline or a small number of anatomical features (Fig. 1n, Extended Data Fig. 2). This feature-based approach accommodated differences between the preparation workflows: the nanoindented surface was measured fresh and hydrated before fixation and staining, whereas the opposing tissue was cryopreserved, cryosection at 10 µm, fixed and stained, and processed using CytAssist. Vascular structures intersecting the original cut plane remained identifiable on both faces, whereas the tissue outline could be altered by processing.

Orientation and scale were adjusted independently for each tissue. The reconstructed grids provided a further internal check on transformation scale. Although the nominal 50 µm grid spacing was not supplied directly to the reconstruction code, it was recovered independently for each ROI from the annotated grid corners. Recorded spacing ranged from 30.4 to 51.0 µm among tissues, corresponding to 61-102% nominal spacing. Among ROIs fitted independently within the same tissue, recovered spacing differed by 0.5-2.6 µm, supporting the internal consistency of grid reconstruction. The tissue-specific change in reconstructed grid spacing agreed with the scaling applied during vascular-feature matching to within 2.1%, providing a computational consistency check. Differences between tissues likely reflect variable deformation during freezing, fixation, and cryosectioning, for which a single global scaling factor would be inappropriate. Some regions visible in the post-nanoindentation preparation were not represented in the final spatial transcriptomic dataset because the thick post-nanoindentation preparation and the 10 µm cryosection sampled different optical volumes, cryosectioning and mounting could deform tissue, and CytAssist-mediated transfer was restricted to the defined capture area. Of the 2,975 candidate stiffness grid points within the co-registered nanoindentation ROIs, 2,771 (93%) containing at least five matched transcriptome bins were retained for analysis. Thus, matched transcriptomic data were available for most candidate co-registered grid points.

Two further checks assessed spatial continuity within the mechanical maps and coarse structural correspondence between the paired surfaces. Stiffness at each indentation grid point correlated with the mean stiffness of its four nearest neighbors, which were separated by 30-51 µm depending on the tissue (pooled Spearman ρ = +0.753; individual ovary ρ = +0.396 to +0.672), indicating spatial continuity within the stiffness maps. Following co-registration, stiffness differed between transcriptomically annotated compartments in six of the eight ovaries tested individually, as described below, supporting coarse correspondence between the mechanical maps and tissue architecture. Together, these analyses assess internal reconstruction and broad anatomical correspondence but do not quantify pointwise registration error between the two surfaces.

### Nanoindentation maps reveal stiffness heterogeneity across fresh ovarian tissue

The ovary is a spatially heterogeneous tissue, with specific structures including corpus luteum, stroma, vasculature, and follicles (Fig. 2a). In order to quantify stiffness across these structures, nanoindentation maps were acquired over distinct regions of the tissue surface. In total, 21 ROIs were acquired and Young’s moduli were determined from the first 2 µm of the loading curve (Extended Data Fig. 3a,b, Supplementary Methods S2 and S3). Aged ovaries were 1.59-fold stiffer than young, but this difference was not significant across the matched cohort of eight ovaries from six mice (95% CI 0.54 to 4.70-fold, *P* = 0.40; Fig. 2b). However, the regional maps showed that stiffness values varied several-fold among adjacent positions within each ovary, demonstrating mechanical heterogeneity that would be obscured by a whole organ average (Fig. 2c,d).

**Figure 2.**
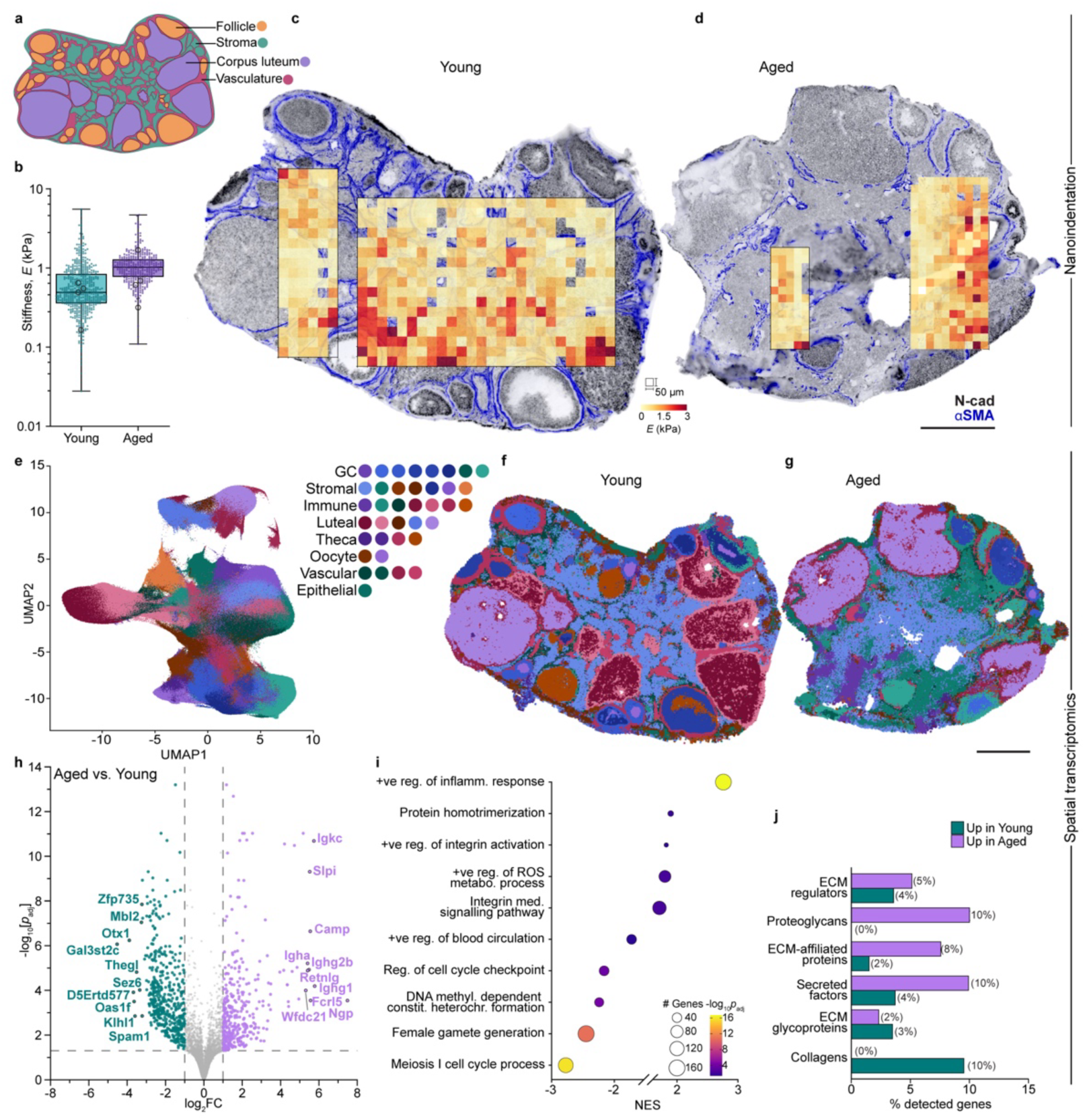
Mechanical and transcriptional organization of young and aged ovaries. (**a**) Four ovarian anatomical compartments used for spatial analysis: corpus luteum, stroma, vasculature, and follicle. (**b**) Stiffness (Young’s modulus, *E*)l, in young and aged ovaries. Box and whisker plots show the median and interquartile range, with whiskers min to max; filled points represent individual 50 µm nanoindentation grid points, and open circles represent per-ovary geometric means. Statistical comparison used a linear mixed model on log_10_transformed stiffness, with animal, ovary, and ROI included as nested random effects. Aged versus young, 1.59-fold, 95% CI, 0.54-4.70, *P* = 0.40. The analysis included 1,648 indentations from 4 young ovaries and 1,123 indentations from 4 aged ovaries, representing 6 mice. (**c,d**) Representative stiffness maps overlaid on IF images from the opposing cryosection of a young (**c**) and aged (**d**) ovary. N-cadherin, grey; αSMA, blue. Nanoindentation grid spacing, 50 µm; Young’s modulus scale 0-3 kPa. Scale bar: 500 µm. (**e**) UMAP of all 4 µm spatial transcriptome bins after BANKSY neighborhood augmentation and Harmony batch correction, colored by 38 annotated cell populations categorized into granulosa, stromal, immune, luteal, theca, oocyte, vascular and epithelial classes. (**f,g**) Spatial distributions of annotated cell identities in representative young (**f**) and aged (**g**) ovaries, colored as in (e). Scale bar: 500 µm. (**h**) Animal-level pseudobulk differential expression between aged and young ovaries. Counts from both ovaries were summed where an animal contributed a pair, yielding on profile per animal. The analysis included n=5 animals per age group, representing 16 ovaries across four Visium HD capture areas Differentially expressed genes are colored teal for upregulated in young, and purple for upregulated in aged. Thresholds denote |log_2_FC| > 1 and Benjamini-Hochberg *P_adj_* < 0.05; genes not meeting both thresholds are shown in grey. (**i**) Gene set enrichment analysis of the ranked differential expression statistic. Dot size represents the leading-edge gene count, and color represents -log_10_(adjusted *P*). Positive NES indicates enrichment in aged tissue. Gene sets comprise Hallmark, GO Biological Process, Cellular Component and Molecular Function, and Reactome collections. Redundant sets were collapsed by leading-edge overlap, and representative terms are displayed. (**j**) Percentage of detected genes in each matrisome category that were differentially expressed (P_adj_ < 0.05, |log2FC| > 1), separated by direction of change. Values indicate detected genes per category: collagens 42/44, ECM glycoproteins 172/194, proteoglycans 30/36, ECM regulators 195/304, ECM-affiliated proteins 132/165, secreted factors 242/367.

Compartments differences were detected only within young tissue. Follicles were stiffer than stroma (1.59-fold, Holm-corrected *P* = 0.0002), corpus luteum (1.38-fold, *P* = 0.008), and vasculature (1.33-fold, *P* = 0.046). No additional within-age comparisons or between age compartment comparisons reached statistical significance (Extended Data Fig. 3c). However, it should be noted that compartment sampling was uneven: corpora lutea and follicles contributed fewer measurements in aged ovaries, and follicles contributed the fewest measurements overall due to their smaller cross-sectional area.

### High resolution spatial transcriptomics resolves cell-, tissue-, and age-related ovarian organization

Visium HD spatial transcriptomic data were evaluated at 2, 4 and 8 µm bin sizes (Extended Data Fig. 4, Extended Data Table 1, Supplementary Methods S4). The 4 µm bins size was selected for downstream analysis because it provided a practical balance between transcript capture and structural detail. Unsupervised clustering after BANKSY neighborhood augmentation and Harmony batch correction resolved 38 distinct transcriptional populations spanning follicular, luteal, stromal, vascular and immune domains (Fig. 2e-g). These populations were annotated using literature-supported marker genes together with their spatial context (Extended Data Fig. 5a,b, Extended Data Table 2). For example, a luteal structure containing infiltrating neutrophils was identified as an early corpus luteum (32), whereas one expressing *Cdkn1a*, *Spp1* and *Cemip* was identified as a regressing corpus luteum. Further, a follicle whose granulosa cells express *Foxo1*, *Ghr* and *Pik3ip1* was identified as atretic. A population expressing the immune markers *Cd79a* and *Ms4a1* localized to stromal aggregates found only in aged ovaries, consistent with previously reported tertiary lymphoid structures (33,34) and was annotated as “Stromal TLS”. Annotations therefore incorporated both transcriptional identity and anatomical location (Supplementary Methods S5). The proportions of annotated bins also shifted with age: luteal identities reduced from ∼50% to ∼15% and follicle-lineage identities from ∼25% to ∼14%, while stromal identities rose from ∼15% to ∼55% and immune identities from ∼4% to ∼12% (Fig. 2f,g, Extended Data Fig. 5c).

Pseudobulk differential expression analysis identified 835 significantly differentially expressed genes associated with age, 353 enriched in aged and 482 in young tissue, with inflammatory and immune programs enriched in aged tissues and gametogenic and cell-cycle pathways enriched in young tissues (Fig. 2h,i, Extended Data Fig. 3d, Extended Data Tables 3 and 4). Specifically, aged tissue was enriched for immune signaling adaptors (*Sit1*, *Lat*, *Grap2)*, immunoglobulin transcripts (*Jchain*, *Igkc*, *Ighg1),* and clustered protocadherins mediating cell adhesion (*Pcdha5*, *Pcdha6*, *Pcdha8* and more), while young tissue was enriched for oocyte and meiotic genes (*Dazl*, *Sycp3*) and MAPK phosphatases (*Dusp14*, *Dusp8*).

Compartment-specific pseudobulk analyses were then used to examine age-associated changes in gene expression and pathways differences within ovarian structures (Extended Data Fig. 6a). Among age-associated ECM genes annotated using the Matrisome Database (35), collagen genes occurred only among transcripts enriched in young tissue, while proteoglycan genes occurred only among those enriched in aged tissue. Other matrisome categories contained genes enriched in both directions, with ECM glycoproteins more frequently enriched in young tissue and ECM regulators, ECM-affiliated proteins, and secreted factors more frequently enriched in aged tissue (Fig. 2j). The collagens driving this enrichment were *Col9a2*, *Col26a1*, *Col11a1* and *Col4a3*, whereas the fibrillar collagens *Col1a1*, *Col1a2* and *Col3a1* did not differ with age. These are transcript-level differences and do not establish corresponding differences in matrix abundance or turnover. Matrisome genes were also cross-referenced against a curated list of mechanobiology and ovary-specific genes to aid interpretation of the subsequent stiffness-gene expression analyses (Extended Data Tables 5 and 6).

### Stiffness-gene expression correlations differ with age on a global tissue level

We next related gene expression aggregated at each 50 µm nanoindentation grid point to its local Young’s modulus across all samples, while adjusting for sequencing depth and age, to identify genes associated with local stiffness. Gene set enrichment analysis was performed on genes ranked by their Spearman correlation coefficients (Fig. 3a, Extended Data Table 8). This pooled analysis identified 594 genes as stiffness-correlated, comprising 487 positive and 107 negative correlations. Because grid point measurements within each ovary were spatially structured and non-independent, we further evaluated the robustness of the gene rankings using a leave-one-ovary-out sensitivity analysis and a stratified analysis of within-ovary associations that controlled for ovary identity (Supplementary Methods S10). Gene-level *P* values were corrected for multiple testing using the Benjamini-Hochberg procedure, with FDR < 0.05 and |ρ| ≥ 0.1 were used to classify stiffness-gene expression associations.

**Figure 3.**
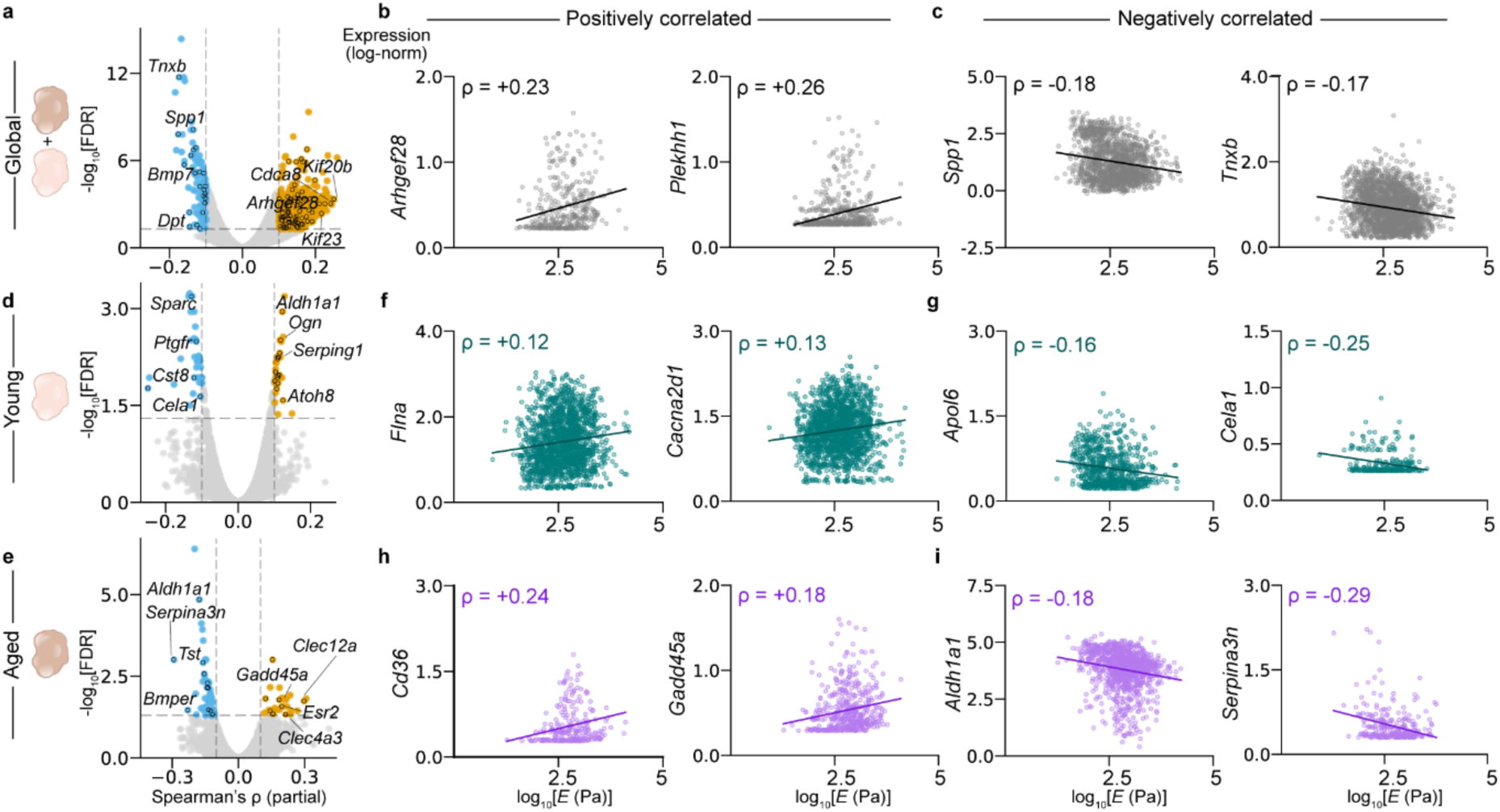
Tissue-level stiffness-expression associations in young and aged ovarian tissue. (**a**) Partial Spearman correlation (ρ) between gene expression aggregated at each 50 µm nanoindentation grid point and the Young’s modulus measured at the same position. The pooled analysis includes 2,771 grid points from eight ovaries of six mice, with sequencing depth and age held constant. Correlations are plotted as −log_10_(Benjamini-Hochberg FDR) against ρ. Dashed lines denote FDR = 0.05 and |ρ| = 0.1. Blue indicates negative correlations meeting both thresholds, yellow indicates positive correlations meeting both thresholds, and grey indicates genes not meeting both thresholds. (**b,c**) Aggregated expression at each grid point plotted against log_10_-transofrmed Young’s modulus for representative positive associations, *Arhgef28* (ρ = +0.23) and *Plekhh1* (ρ = +0.26) (**b**), and negative associations, *Spp1* (ρ = −0.18) and *Tnxb* (ρ = −0.17) (**c**). Each point represents one 50 µm grid point. Lines show unadjusted linear fits for visualization, whereas the reported ρ values are partial Spearman correlations. (**d,e**) The same analysis within young tissue, comprising 1,648 grid points from four ovaries (**d**), and aged tissue, comprising 1,123 grid points from four ovaries (**e**). Correlations in (**d**-**i**) were calculated separately within each age group, with sequencing depth and ovary identity held constant. (**f**) Representatively positive association in young tissue: *Flna* (ρ = +0.12) and *Cacna2d1* (ρ = +0.13). (**g**) Representative negative associations in young tissue: *Apol6* (ρ = −0.16) and *Cela1* (ρ = −0.25). (**h**) Representative positive association in aged tissue: *Cd36* (ρ = +0.24) and *Gadd45a* (ρ = +0.18). (**i**) Representative negative associations in aged tissue: *Aldh1a1* (ρ = −0.18) and *Serpina3n* (ρ = −0.29).

The 487 genes whose expressed increased with stiffness (i.e., ‘positively correlated’) were enriched for cytokinesis and chromosome segregation programs (Fig. 3a). Among the individual genes positivity correlated with stiffness were the mitotic kinesins *Kif20b* (ρ = +0.25) and *Kif23* (ρ = +0.22) and the chromosomal passenger component Cdca8 (ρ = +0.24). Representative positive correlations included the Rho guanine nucleotide exchange factor *Arhgef28* (ρ = +0.23) and *Plekhh1* (ρ = +0.26; Fig. 3b). The 107 genes whose expression increased as stiffness decreased (i.e., ‘negatively correlated’) were enriched for mitochondrial respiratory electron transport, cholesterol biosynthesis, and sterol biosynthesis. Representative genes negatively correlated with stiffness included the matrisome-associated genes *Spp1* (ρ = -0.18), *Tnxb* (ρ = -0.17), *Bmp7* (ρ = -0.16) and *Dpt* (ρ = -0.15; Fig. 3c).

We then carried out the same analysis by age group, which resulted in differences in both the number and composition of the ranked genes. Specifically, 73 genes were identified to be correlated to stiffness in young ovaries (Fig. 3d) and 84 genes in aged, (Fig. 3e). In young tissue, representative positive associations with stiffness included the actin crosslinker *Flna* and voltage-dependent calcium channel genes *Cacna2d1* (Fig. 3f), while the apolipoprotein gene *Apol6* and elastase gene *Cela1* were negatively associated (Fig. 3g). In aged tissue, the scavenger receptor *Cd36* and stress-responsive gene *Gadd45a* were positively associated, while the retinoid enzyme *Aldh1a1* and serine protease inhibitor *Serpina3n* were negatively associated (Fig. 3i). Comparing the two age groups directly, 146 genes met the prespecified threshold for a difference in correlation strength between age groups, with 81 in young and 65 in aged tissue (Extended Data Fig. 6b). Correlation coefficients shifted toward more positive values with age for immune regulators, including *Cited1* (Δρ = +0.54), *Rasgrp1* (+0.45), and *Ptpn22* (+0.37), and toward more negative for matrix and stromal genes, including *Serpina3n* (Δρ = -0.36), *Dpt* (-0.30) and *Aldh1a1* (-0.30; Extended Data Fig. 6b).

### Compartment-resolved analyses identify context-specific stiffness associations

We next assigned the 4 µm spatial transcriptome bins and matched nanoindentation grid points to the four major ovarian compartments: corpus luteum (CL), stroma (S), vasculature (V), follicle (F) (Fig. 4a, b, Extended Data Figs. 7 and 8). Within each ROI, compartment identity, matched stiffness, and spatial gene expression could then be examined in the same coordinate space (Fig. 4b-d). Restricting the stiffness-gene expression correlation analysis to individual compartments returned 781 associations across 733 genes, compared to 594 genes in the whole-tissue analysis.

**Figure 4.**
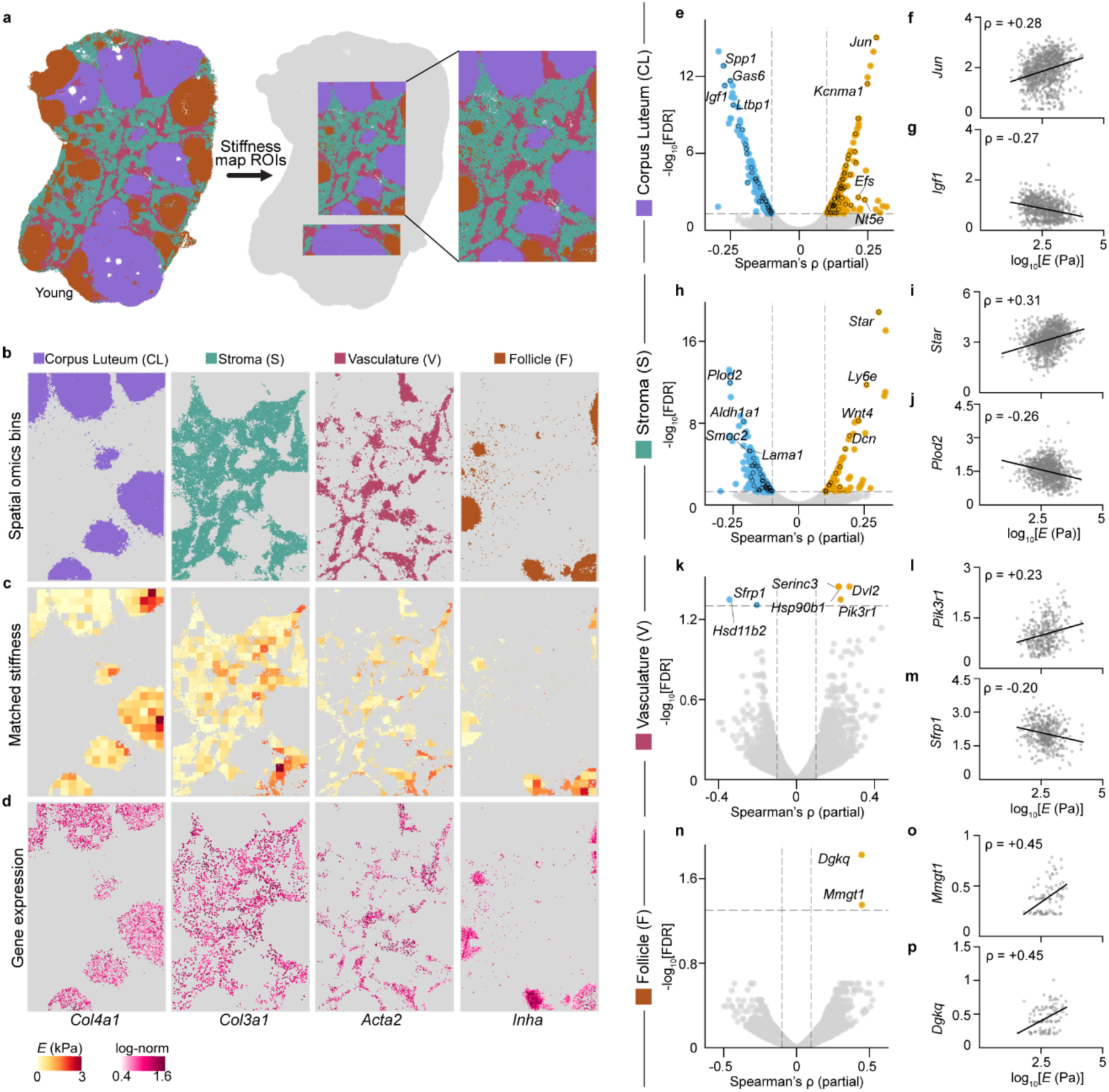
Compartment-resolved local stiffness-expression associations. (**a**) Representative spatial transcriptome data of a young ovary colored by compartment: corpus luteum (CL, purple), stroma (S, teal), vasculature (V, pink), follicle (F, brown), with matched nanoindentation ROIs outlined. (**b**) Assigned 4 µm spatial transcriptome bins within the ROI, separated by compartment. (**c**) Stiffness at the matched nanoindentation 50 µm grid points within each compartment. (**d**) Spatial expression of representative compartment-associated genes within the ROI: *Col4a1* in CL, *Col3a1* in S, *Acta2* in V and *Inha* in F. (**e-p**) Compartment-resolved partial Spearman correlations (ρ) between aggregated gene expression and Young’s modulus, with sequencing depth and age held constant. Each measurement represents one 50 µm nanoindentation grid point. CL included 1,037 measurements (**e**), with representative positive and negative associations for *Jun* (ρ = +0.28) (**f**) and *Igf1* (ρ = -0.27) (**g**), respectively. S included 1,040 measurements (**h**), with representative associations for *Star* (ρ = +0.31) (**i**) and *Plod2* (ρ = −0.26) (**j**). V included 435 measurements (**k**), with representative associations for *Pik3r1* (ρ = +0.23) (**l**) and *Sfrp1* (ρ = −0.20) (**m**). F included 148 measurements (**n**), with positive associations for *Mmgt1* (ρ = +0.45) (**o**) and *Dgkq* (ρ = +0.45) (**p**). Lines in the expression plots show unadjusted linear fits for visualization; reported ρ values are partial correlations.

Corpus luteum contributed 424 stiffness-correlated genes (Fig. 4e), including positive associations for the immediate-early transcription factor *Jun* (Fig. 4f) and the collagen-modifying protein *Colgalt2,* and negative associations for the metalloendopeptidase *Mmel1* and insulin-like growth factor 1 gene *Igf1* (Fig. 4g). Stroma contriuted 349 stiffness-correlated genes (Fig. 4h), including positive correlations for steroidogenic gene *Star* (Fig. 4i) and the Wnt ligand *Wnt4*, and negative correlations for the ECM-modifying genes *Cav1* and *Plod2* (Fig. 4j). Vasculature contributed six stiffness-associated genes (Fig. 4k), including positive associations for the Wnt pathway regulator *Dvl2* and PI3K regulator *Pik3r1* (Fig. 4l), and negative associations for the Wnt regulator *Sfrp1* (Fig. 4m) and hormone-metabolizing enzyme *Hsd11b2*. Follicles contributed only two positively correlated genes to stiffness (Fig. 4n), namely the membrane transporter *Mmgt1* (Fig. 4o) and steroidogenic-signaling lipid kinase *Dgkq* (Fig. 4p). Of the 733 compartment-correlated genes, 611 genes (83%), did not appear in the whole-tissue tests (Fig. 3), indicating that compartment-resolved analysis recovers associations for genes relevant to each compartment’s own biology that are obscured by pooling across the tissue structures (Extended Data Table 6).

Interpreatation of these compartment-level associations requires accounting for uneven sampling across ages and ovaries. Of the 1,037 corpus luteum measurements, 966 came from young ovaries, while 739 of the 1,040 stromal measurements came from aged ovaries. The pooled compartment results may therefore reflect variation between ages and ovaries in addition to variation among positions within an ovary. Further, follicles occupied only ∼5.3% of the nanoindentation grid points, corresponding to 148 matched measurements, and yielded two stiffness-correlated genes (Fig. 4n). This result reflects limited coverage rather than evidence against follicular mechanosensitivity. Recovering a compartment-specific signature within an ovary therefore requires adequate representation of each compartment in multiple ovaries at both ages.

Applying a stricter within-ovary test across age retained 31 of the 424 corpus luteum associations, one of the 349 stroma associations, one of six vasculature associations, and none of the follicular associations (Extended Data Fig. 6c, Extended Data Table 6). The retained corpus luteum associations formed a coherent lipid-metabolism signature. Seven cholesterol biosynthesis enzymes (*Sqle*, *Cyp51*, *Msmo1*, *Idi1*, *Fdps*, *Hmgcr*, and *Tm7sf2*), the SREBP regulator *Insig1*, the high-density lipoprotein receptor *Scarb1* and the lipogenic enzymes *Acly*, *Scd1* and *Me1* decreased as local stiffness increased. By contrast, the steroidogenic transcription factor *Nr5a1* and the collagen receptor tyrosine kinase *Ddr2* increased with stiffness. Thus, inverse associations with cholesterol and sterol biosynthesis were among the few compartment-level patterns retained after adjustment for ovary identity. The remaining associations were also metabolic: the lipogenic transcription factor *Mlxipl* in stroma (ρ = +0.30) and the steroid-metabolizing enzyme *Hsd11b2* in vasculature (ρ = -0.39).

## Discussion

We developed a paired-surface spatial mechanomics workflow that places nanoindentation maps acquired on fresh tissues and 10x Genomics Visium HD spatial transcriptomic profiles from the immediately adjacent, parallel tissue surface into a common coordinate frame. Across 21 independently aligned nanoindentation map regions, the workflow produced 2,771 matched stiffness-gene expression measurements with a median of 89 4-µm transcriptome bins assigned to each 50 µm nanoindentation grid point. The shared α SMA-positive vascular features provided distributed landmarks for setting tissue orientation and scale. Agreement in the recovered grid spacing among independently reconstructed ROIs of the same tissue to within 2.6 µm, spatial continuity between neighboring stiffness measurements (pooled ρ = +0.75, >+0.3 in all eight ovaries), and compartment-level stiffness differences in six of eight ovaries provided complementary checks of grid reconstruction and coarse anatomical correspondence. These controls do not directly quantify pointwise registration error between surfaces. Future incorporation of quantitative landmark validation and deformable image registration, as implemented in common coordinated and diffeomorphic alignment approaches for spatial transcriptomics (36,37), could improve the local correspondence. The workflow is designed to be accessible to laboratories with nanoindentation and spatial omics capabilities, using a landmark-based alignment that can be performed in spatial viewers with graphical alignment tools.

Application to the aging ovary, which exhibits high spatial heterogeneity and dense interstitial ECM structures, demonstrates how direct mechanical measurements provide information not captured by spatial molecular profiling. We found that compartment composition alone was only weakly associated with local stiffness, with the strongest association across the 38 annotated cell populations and four anatomical compartments reaching ρ = +0.056 (*P* = 0.003, FDR = 0.045, n = 2,771 grid points). This weak relationship is consistent with contributions from fibrillar ECM components, matrix architecture, and poroelastic fluid movement that are not represented by compartment identity alone (38). In pooled analyses, softer regions expressed more matrisome genes, with ECM glycoproteins enriched 9.7-fold among genes negatively correlated with stiffness, alongside mitochondrial respiratory electron transport and cholesterol biosynthesis. By contrast, stiffer regions expressed more cytokinesis and chromosome segregation programs. Reduced expression of matrix-associated genes does not establish reduced protein abundance or explain the measured Young’s modulus. Instead, the inverse association links local mechanical state with transcriptional programs involved in matrix homeostasis. An *in vitro* study similarly showed that reducing compressive stress on isolated follicles increased *Fbln5*, *Fbn1, Col4a1*, *Col4a2*, and *Col4a3* expression in granulosa cells, which was interpreted as a compensatory matrix response (21).

Collagen provides a related example of why mechanical and molecular states should be measured together. Collagen transcripts were enriched in young ovaries in the age-based comparison (Fig. 2j), although this pattern was driven by basement membrane and fibril-associated collagens (*Col4a3*, *Col9a2*, *Col11a1*, *Col26a1*) rather than the fibrillar collagens *Col1a1*, *Col1a2,* and *Col3a1*, which did not differ with age. Matrix-related genes were also enriched among negative stiffness correlations (Fig. 3a,c), despite previous reports of increased ovarian fibrosis and stiffness with age (3,6,7,20). These observations reinforce that transcript abundance is not a direct proxy for matrix abundance, organization, or mechanics. Long protein lifetimes, altered turnover, crosslinking, and matrix architecture could all contribute to this apparent discordance, but the present data cannot distinguish these possibilities, nor determine whether these transcriptional programs cause, respond to, or merely co-localize with the measured mechanical state. Spatial measurements of protein abundance, matrix architecture, and composition will be required to distinguish these possibilities.

Young’s modulus was estimated to be 1.59-fold higher in aged ovaries, but was not significant. The nanoindentation maps instead reflect the spatial mechanical heterogeneity within and between ovaries. Per-ovary geometric means spanned a 4.1-fold range in young ovaries and a 5.5-fold range in aged ovaries, while variance between ovaries exceeded variance between animals in the compartment model. However, the small cohort, unstaged estrous cycles, and unequal compartment representation prevent these observations from establishing greater mechanical heterogeneity or increased stiffness with age. Thus, a larger, estrous-staged cohort will be required to estimate age-associated changes in spatial ovarian mechanics.

Age altered the identities of stiffness-associated genes more clearly than their number. Similar numbers of genes were ranked as stiffness-associated in young and aged ovaries, while 146 genes met the prespecified threshold for a difference in correlation strength between age groups. Mitochondrial oxidative phosphorylation declined with increasing stiffness at both ages, cholesterol and sterol biosynthesis declined with stiffness in young tissue, and phagocytosis, interleukin-6 production, and reactive oxygen species metabolism increased with stiffness in aged tissue. The aged inflammatory signature included macrophage identity genes and may therefore reflect enrichment of myeloid cells in stiffer regions rather than altered transcription within individual cells. Young vasculature was also enriched for Hippo-pathways genes, and the YAP and TAZ cofactor Tead1 correlated with stiffness in young but not aged tissues. Together, these findings indicate age-dependent reorganization of spatial stiffness-gene expression associations rather than a uniform loss of association with age. They do not distinguish contributions from altered cell composition, persistent mechanical exposure, or changes in cellular mechanosensitivity, which will require perturbation and time course experiments.

Compartment-resolved analyses illustrate both the capability and the sampling requirements of the method. Adjustment for ovary identify removed most pooled compartment associations, but the retained corpus luteum genes formed a coherent lipid-metabolism signature. Cholesterol biosynthetic enzymes, the SREBP regulator *Insig1*, the high-density lipoprotein receptor *Scarb1*, and several lipogenic enzymes decreased as local stiffness increased. Because lipoprotein uptake and de novo synthesis provide compensatory cholesterol sources for luteal steroidogenesis (41), their concurrent decline may reflect reduced cholesterol demand during corpus luteum regression rather than direct mechanical regulation. A corpus luteum forms and regresses within days, making its mechanical and metabolic state strongly dependent on the stage sampled. This poses a challenge for aged ovaries, which have fewer follicles and corpora lutea due to follicle depletion and reduced ovulation. Estrous staging would match corpus luteum number and stage within an age group, but it cannot equalize compartment representation between young and aged tissues. The 50 µm nanoindentation grid also provided limited follicle coverage, preventing a conclusive follicle-specific analysis. Follicles were stiffer than stroma, corpus luteum, and vasculature in young tissue, whereas no compartment differences were detected in aged tissue. This differs from some previously reported measurements (7, 22), and may reflect differences in measurement scale, tissue preparation, estrous stage, or compartment sampling. Future studies of compartment-resolved stiffness-gene expression associations should therefore balance or enrich structures of interest across animals and experimental conditions.

Several design features define the interpretation of these measurements. Vibratome slicing releases long-range tissue tension, so intact organ mechanics are not preserved. The two modalities are acquired from adjacent, rather than identical surfaces, and the surfaces undergo different preparation workflows, which can alter tissue geometry and limit correspondence with small structures. Nevertheless, 93% of candidate co-registered grid points met the transcriptome-bin inclusion criterion.

These analyses support the internal consistency of grid reconstruction and coarse anatomical correspondence but do not quantify target registration error at individual measurement positions, which would require corresponding landmarks identified independently on both surfaces and excluded from the registration procedure. Each 50 µm nanoindentation grid point aggregates a median of 89 4-µm spatial transcriptome bins, which improves transcript detection and accommodates small positioning uncertainties but sacrifices cellular specificity. Lastly, our ROIs predominantly sample the ovarian interior, which means the stiffer cortical surface is not sampled, where AFM has shown localized increased stiffness with age and in disease (3,5).

Within these constraints, paired-surface spatial mechanomics provides a strategy for relating experimentally measured tissue mechanics to spatial molecular state at a local tissue scale. The workflow is adaptable to other fresh, sectionable tissues for which mechanical cues play a major role, including tumors, fibrotic scars, and vessel walls (42,43). The approach could be extended through additional omics layers or structural ECM imaging. Imaging-based spatial transcriptomics performed directly on the section could eliminate CytAssist-mediated transfer as one potential source of data loss, although differences between adjacent tissue surfaces would remain. Ultimately, perturbation and longitudinal experiments will be required to establish causal relationships between molecular and mechanical states.

## Methods

### Animals and tissue collection

Ovaries were obtained from reproductively young (arrived at 9 weeks old, studied at 3 months old) and reproductively aged (arrived at 12 months old, studied at 14 months) female CD-1 (ICR) mice sourced from BioLASCO (Taiwan) and housed in the National University of Singapore Comparative Medicine animal facility under pathogen-free conditions. Mice were housed in individually ventilated cages at five per cage with ad libitum access to food and water on a 12 h light and 12 h dark cycle, at 18-25 °C and 30-70% relative humidity. All procedures were approved by the Institutional Animal Care and Use Committee of the National University of Singapore (protocol R22-0511).

Mice were euthanized by carbon dioxide asphyxiation followed by cervical dislocation. Ovaries were harvested into Leibovitz’s L-15 medium (#11415064, Gibco) supplemented with 3 mg/ml bovine serum albumin (#A9418, Sigma), and fat and bursal membranes were removed under a dissecting stereomicroscope. Estrous stage was not determined.

Sixteen ovaries from ten animals were processed for spatial transcriptomics, comprising five animals per age group. Six animals contributed a contralateral pair and four contributed a single ovary. Of these, eight ovaries from six mice carry matched nanoindentation measurements on the adjacent tissue surface; each age group contained three mice, one of which contributed both ovaries. The animal was the experimental unit for age-based inference. Counts from contralateral ovaries were summed within each animal for pseudobulk differential expression, whereas ovary, ROI, and grid-point identifiers were retained for the spatial mechanical analyses. Animal, ovary, section, and ROI identifiers for every sample are provided in Extended Data Table 7.

### Vibratome sectioning and paired-surface preparation

Each ovary was embedded in 3% (w/v) low-melting-point agarose (16520100, Invitrogen) in sterile 1X PBS (Phosphate-Buffered Saline Solution, Gibco) in a cryomold and set at 4°C for 5 min. The agarose block was trimmed and glued to a precooled vibratome stage with cyanoacrylate adhesive, and mounted in a bath of ice-cold 1X PBS on a Leica VT1200S vibratome maintained on ice and fitted with a fresh Feather blade (121–9) for each ovary. Trimming was performed at 0.05 mm/s, amplitude 1 mm and autofeed 500 µm; sectioning was performed at 0.02 mm/s, amplitude 1 mm and autofeed 100 µm, to produce 100 µm slices (Fig. 1c). All sectioning was completed within 1 h of euthanasia. Sections were kept in L-15 medium at 4°C until indentation, and the second section of the contralateral ovary was indented directly after the first was complete.

Slices were attached to poly-D-lysine-coated slides and immersed in L-15 medium within a 3D-printed chamber sealed with duplicating silicone eco-sil speed (#13007100, Picodent Twinsil). Attachment was verified by topping up medium over the section. A section that remained in place rather than lifting under its own buoyancy was taken as attached. A section that failed to attach was discarded and a further section taken onto a fresh coated slide. Slide cleaning, coating, and chamber specifications are given in Supplementary Methods S1.

After attachment was confirmed, the remaining tissue was sectioned was sectioned to produce a 1 mm thick block. The nanoindented face of the removed 100 µm section (surface 1) and the upper face of the remaining block (surface 2) were the two opposing faces generated by the same vibratome cut. Surface was therefore immediately adjacent to surface 1 and was cryopreserved for special transcriptomics. This parallel geometry is what establishes correspondence between the two modalities.

### Nanoindentation

Nanoindentation was performed on the attached sections under L-15 medium at room temperature with an Optics11Life Chiaro nanoindenter using a spherical probe (spring constant *k* ≈ 0.025 N m^-1^; tip radius *R_tip_* ≈ 27.5 µm). Before each experiment, the probe spring constant was calibrated in L-15 medium according to the manufacturer’s protocol through the geometry factor and conversion of cantilever deflection to applied force. Nanoindentation measurements were performed in displacement control mode with a 0.5 s baseline hold, loading to a 10 µm piezo displacement over 2 s (5,000 nm/s), a 1 s dwell at peak, retraction over 2 s (5,000 nm/s), a 0.5 s post-retraction hold, and a sampling rate of 1,000 Hz. Measurements were acquired on rectangular grids at 50 µm center-to-center spacing. Each measurement position is hereafter referred to as a grid point, and each grid defines an ROI. ROIs were positioned within the tissue interior rather than at the tissue margin to protect the probe, preferentially sampling the medulla and inner cortex; the outer cortex is therefore under-represented. A total of twenty-one ROIs were acquired across the eight ovaries. Probe start and end positions were recorded by inverted microscope for subsequent co-registration. Nanoindentation on the first ovary began within 1 h of euthanasia and was completed within 4-6 h for each ovary. The second ovary was maintained in L-15 media during the first ovary indentation.

### Nanoindentation data processing and quality control

Force-displacement curves were processed with a custom Python pipeline (Supplementary Code 1) adapted from the atomic force microscopy analysis code of Biswas et al. (21) and modified for nanoindentation input data, with quality control (QC) and per-slice summaries generated by Supplementary Code 2 to 4, stiffness distributions and their statistical comparison by Supplementary Code 5, and stiffness maps by Supplementary Code 6. The Young’s modulus, *E*, was estimated from the first 2 µm of the loading curve using the Hertz contact model for a spherical indenter,

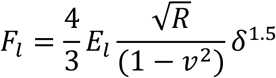

where *F_l_* is the loading force, *δ* is indentation depth, *R* is the effective probe radius, and *v* = 0.49 is the assumed Poisson’s ratio for a nearly incompressible tissue. Because the tissue surface was locally flat relative to the probe, R was taken as R_tip_. The fitted Young’s modulus, *E*, was the primary mechanical parameter and is referred to as stiffness throughout. Ovarian tissue is viscoelastic, so *E* is considered an apparent modulus conditional on the Hertz contact model, the assumed Poisson’s ratio and the stated loading conditions. Comparisons are therefore interpreted between measurements acquired under identical settings rather than as an assumption-free material constant.

Quality control was applied to the data in the following order: (1) curves reporting a maximum load of −0.001 µN were excluded as failed indentations; (2) curves with surface contact model residual sum of squares ratio above 1.655 were retained; (3) fits with a relative amplitude standard error >0.15 were excluded; and (4) fitted moduli outside 7.6 to 22,726 Pa were excluded as the limits of the measurable range for this probe as determined from its calibrated spring constant and tip radius.

Of 5,049 acquired curves across 8 ovaries contributing to the reported cohort, 855 (16.9%) failed contact detection, 150 (3.0%) made contact but did not return a fitted modulus, and 298 (5.9%) failed one or more fit quality or modulus range criteria. In total, 3,746 (74.2%) measurements passed the QC. Of these, 2,975 were within the spatial transcriptomics-eligible ROIs, and 2,771 were assigned at least five 4 µm transcriptome bins and entered the stiffness-gene expression analysis. Threshold derivation and sensitivity are given in Supplementary Methods S3.

### Post-nanoindentation tissue preservation, immunofluorescence, and confocal imaging

Indented sections remained attached to the marked glass slide, which was placed in a sealed petri dish and stored at -80°C for storage until staining. Maintaining the section on the same marked slide preserved its position and orientation relative to the recorded probe locations, allowing the nanoindentation grid to be referenced during image alignment. Sections were later fixed and stained as a single batch using the protocol described below for the paired cryosections, using identical fixation, permeabilization, methanol treatment, and antibody panel and concentrations, allowing αSMA-positive vasculature was resolved on both faces of the vibratome cut. Sections were imaged on a BC43 benchtop spinning disk confocal microscope (Andor, Oxford Instruments) with a 20x, N.A. 0.8 objective, acquiring z-stacks at 0.982 µm intervals to a depth of 50-60 µm from the indented surface. The local tissue surface was identified across the x-y field using the shallowest optical sections containing Hoechst-positive nuclei. For sections exhibiting approximately planar tilt relative to the imaging plane, the spatially calibrated image volume was resliced into an orthogonal view in Fiji, and the angle of the superficial tissue boundary was measured using the line tool. The volume was rotated by this angle using bilinear interpolation and resliced such that the digitally reconstructed superficial tissue plane was parallel to the imaging plane (Supplementary Code 18). The same transformation was applied to all fluorescence channels. Maximum intensity projections were generated over the first 12 µm below the locally defined surface using Imaris (v.10.2.0) and used for registration. The depth was selected to approximate the superficial volume represented by a paired 10 µm cryosection while retaining sufficient αSMA-positive vasculature signal for feature matching. During co-registration in SpatialX, rotation and scaling were determined independently for each tissue by matching vascular features; no global tissue shrinkage factor was assumed.

### Cryosectioning, immunofluorescence, and Visium HD capture

The paired 1-mm block was embedded in Tissue-Tek^®^ O.C.T. (#4583, Sakura Finetek USA) with the surface 2 facing up and parallel to the base of the mold, and fresh frozen by lowering the cryomold into isopentane chilled in dry ice, following the 10x Genomics fresh-frozen tissue preparation protocol (CG000763 Rev E). Blocks were stored in sealed containers at -80°C for 6.5 to 17.4 months before cryosectioning in a single batch (Fig.1h). Storage duration per ovary is given in Extended Data Table 7. Parallelism between the block face and the cutting plane was verified during sectioning, and each section was inspected before collection to confirm that surface 2 had been reached (Supplementary Methods S1). Blocks were equilibrated to -20°C for at least 30 minutes and cryosectioned at 10 µm in one processing batch with the blade at -20°C and the specimen head at -10°C. Two to three serial 10 µm sections were collected per ovary, beginning directly from surface 2. Before data acquisition, one section was selected for Visium HD processing using the prespecified criteria of complete tissue transfer, placement within the capture area, and absence of visible section damage.

IF was performed on tissue sections following the 10x Genomics fresh-frozen protocol (CG000763 Rev E), which minimizes the time between fixation, staining, and imaging to limit mRNA degradation. Briefly, sections were fixed in 4% paraformaldehyde (#157-4, Electron Microscopy Sciences) in 1X PBS for 30 min at room temperature, washed in 1X PBS, permeabilized in 1% sodium dodecyl sulfate for 2 min with 1X PBS washes on either side, and incubated in pre-chilled 70% methanol for 60 min on ice, before staining with conjugated antibodies against αSMA conjugated with AlexaFluor 488 (#53-9760-80, Invitrogen; 2.5 µg/ml), N-cadherin conjugated with CoraLite 594 (#CL594-22018, ProteinTech; 5 µg/ml), and Hoechst 33342 (#H3570, Invitrogen; 0.2 µg/ml) over 1 h at room temperature in the dark, and then imaged on a Zeiss Axioscan 7 with a 20x (N.A. 0.8) objective using identical exposure settings for both Visium HD slides, acquiring multi-plane Z-stacks before merging them using the Extended Depth of Focus (EDF) function to obtain a fully in-focused 2D image.

After imaging, sections were transferred with a Visium CytAssist ( CAVG10400, software version v2.1.0.14, 10x Genomics) onto Visium HD slides (#PN-1000675, 10x Genomics) within the capture area (6.5 mm x 6.5 mm) and processed for probe hybridization, ligation, release, and library construction according to the manufacturer’s protocol for Visium HD Mouse Transcriptome (#PN-1000676, 10x Genomics). Libraries were sequenced on the Illumina NovaSeq X Plus 25B flow cell, yielding 475 to 609 million reads per capture area. Library size distributions and concentrations for each capture area are provided as Supplementary File 1. A total of four capture areas (CA_01 to CA_04) were used; the sample layout and sequencing metrics are in Extended Data Table 1. Space Ranger web summaries for each capture area at 8 µm binning are provided as Supplementary File 2.

### Spatial transcriptomics data processing and integration

Tissue alignment was performed on Loupe Browser v8.1.2 using CytAssist images of the fiducial markers on the slides to produce alignment JSON files. FASTQ files, the Visium mouse transcriptome probe set v2.0 for mm10 CSV file, CytAssist image data, IF image data, and alignment JSON files were processed on Space Ranger v3.1.3 for 2, 4, and 8 µm bin resolutions, aligning to mm10 (GRCm38) mouse reference genome. Filtered feature-barcode matrices from the four capture areas were loaded as on-disk sparse matrices with BPCells v0.3.1 and processed in R v4.5.1 using Seurat v5.3.1 (44). Bin barcodes were made unique by appending capture-area identifiers, and Ensembl IDs converted to gene symbols using Space Ranger annotations.

A 4 µm bin size was used for all downstream analyses. Bin size was selected by comparing per-bin UMI counts, per-bin detected features, principal component structure, cluster separability and spatial cluster coherence at 2, 4 and 8 µm (Extended Data Fig. 4, Supplementary Methods S4, Supplementary Code 7). Bins containing fewer than 50 UMIs were excluded, and retained bins were normalized using Seurat’s LogNormalize function using a median library-size scale factor of 271 counts, followed by variable feature selection by variance-stabilizing transformation, retaining the top 2,000 features. BANKSY v1.6.0 (45) was applied to augment each bin’s profile with a spatially weighted average of its six nearest spatial neighbors (k_geom = 6, λ = 0.3, 20 principal components). Batch effects across capture areas were corrected by Harmony v1.2.4 (46) applied to the BANKSY principal components, with capture area identity as the sole batch variable and a maximum of 20 iterations, without supplying age or cell population labels. The resulting embedding was used for UMAP and Louvain clustering over dimensions 1 to 20. Clustering was evaluated across resolutions 0.2 to 1.0 by clustree and resolution 0.8 was selected based on cluster stability across resolutions and concordance with established ovarian marker and spatial patterns, yielding 43 transcriptionally distinct clusters. Integration, clustering and marker gene identification were performed with Supplementary Code 8.

### Transcriptome bin population annotation

Because a 4 µm bin can contain transcripts from portions of one or more cells, annotations describe the dominant transcriptional population represented by each bin rather than a segmented single-cell identity. A 4 µm bin samples at close to single cell scale in the smallest populations while remaining large enough to retain usable transcript counts, which is why identities were assigned at this resolution rather than at coarser binning. Differentially expressed marker genes were identified per cluster by one-versus-all Wilcoxon rank-sum test (log_2_FC ≥ 0.5, adjusted p < 0.05) and ranked by a composite score integrating effect size, detection specificity, and statistical confidence (MG_score = avg_log2FC × (pct.1 − pct.2) × (−log10(p_val_adj + 1×10⁻³⁰⁰))). Bin-level Wilcoxon *P* values were used for marker ranking rather than population-level inference, since neighboring spatial bins are not independent and are further correlated by BANKSY neighborhood augmentation. Annotations integrated cluster-enriched markers, spatial localization, tissue morphology, and αSMA immunofluorescence with a curated published ovarian cell-type reference panel for the female reproductive tract (47) and cross-referencing published mouse and human ovarian single-cell and spatial transcriptomic atlases (23, 29, 48–51). Five clusters comprising fewer than 50 bins were excluded as artefacts because they did not cluster with any other bin populations, leaving 38 annotated populations numbered 0 to 37 (Fig. 2e, Extended Data Fig. 5). Full marker gene evidence, pairwise comparisons and annotation rationale for every population are given in Supplementary Methods S5 (Extended Data Table 2). Clusters 6, 24, and 36 were flagged rather than excluded and were retained for all analyses. The flag and its basis are given per cluster in Supplementary Methods S5.

### Compartment assignment

The 38 annotated populations were grouped into follicle (F), corpus luteum (CL), vasculature (V), immune, stromal (S), and ovarian surface epithelium. For the four-compartment display (F, CL, V, S), immune bins were combined with vascular and ovarian surface epithelium was combined with stroma. Assignment used cell identity as the primary criterion, with spatial boundary overriding it where the two conflicted. ROI and bin assignment was performed with Supplementary Code 9.

An indentation grid point was assigned to a compartment when >50% of its expression-matched 4 µm bins belonged to that compartment; indentations reaching 50% in no compartment were retained for whole-tissue analyses and excluded from compartment analyses, and the small number reaching exactly 50% in two compartments were assigned to both. The number of measurements per ovary per compartment ranged from 4 to 433, and three combinations were absent: follicle in Y5 and A1, and corpus luteum in A2. For a compartment-specific profile, raw counts were summed only over bins of that compartment, and the corresponding compartment-specific total count was used as the library size offset.

Of 3,915,673 bins under tissue areas, 3,728,148 passed the 50-UMI threshold and were assigned, with 688,388 to F, 1,187,901 to CL, 697,614 to V and 1,154,245 to S, comprising 1,797,568 bins from young and 1,930,580 bins from aged tissue. Of these, 269,964 bins fall within nanoindentation ROIs, of which 258,049 carry a matched measurement and 255,620 a valid modulus, and these enter the stiffness-expression analysis.

### Pseudobulk differential expression, gene set enrichment, and matrisome classification

Differential expression was computed on pseudobulk profiles. Raw counts were summed across all bins of a given compartment or annotated population within each animal. Counts from contralateral ovaries were combined when both ovaries were available, yielding one animal-level profile per test and from 5 animals per age group for whole tissue analysis, representing 16 ovaries. Profiles derived from <10 bins were excluded, and genes were retained if they exceeded 5 counts in at least two profiles. No animal fell below the bin threshold in any compartment or population comparison. Testing used DESeq2 v1.50.2 (52) with a Wald test of the ordinary null, log_2_FC = 0. The ordinary null was used throughout, and a threshold-based alternative was computed for reference only (Supplementary Methods S6). Fold-change and significance thresholds (|log_2_FC| > 1, Benjamini-Hochberg adjusted p < 0.05) were applied after testing, for counting, coloring and labelling only. Both directions are reported throughout; negative fold changes denote higher expression in young tissue (Fig. 2h, Extended Data Fig. 6a). Fold-change estimates were shrunk with apeglm v1.32.0 (53) for visualization, with the unshrunken maximum likelihood estimate also reported, since only the latter is the quantity the Wald test evaluated. Differential expression, enrichment and matrisome classification were performed with Supplementary Code 10, and enrichment dot plots curated with Supplementary Code 11.

Gene set enrichment analysis (GSEA) was assessed with fgsea v1.36.2 (54) on the complete ranked gene list, ordered by the signed Wald statistic. Gene sets were drawn from the mouse-native MSigDB release 2026.1.Mm (55,56) with no human-to-mouse ortholog mapping, comprising Hallmark (50 sets), GO Biological Process (7,781), Cellular Component (1,067) and Molecular Function (1,899), and Reactome (1,333), giving 12,130 sets in total. Two redundancy controls were applied. collapsePathways identifies pathways remaining significant after conditioning on a more significant one, which resolves parent-child nesting within GO. Clustering of significant pathways by leading-edge gene overlap identifies pathways drawing their enrichment from the same genes under unrelated names, and is necessary because MSigDB set names reflect the experimental context in which genes were first characterized rather than the tissue under study. Enrichment is reported as leading-edge clusters with their constituent genes rather than as a count of significant pathway names (Fig. 2i, Supplementary Methods S7). Differentially expressed genes were additionally classified against the matrisome database (35) into matrisome categories within the core matrisome and matrisome-associated divisions, and reported as the percentage of detected genes per category changing in each direction (Fig. 2j).

### Image co-registration

Registration comprised four steps, as exemplified in Extended Data Fig. 1 for one tissue sample; vascular-feature matching for all eight ovaries is shown in Extended Data Fig. 2. First, top-view stereoscope and bottom-view inverted microscope images were aligned using physical landmarks, and recorded probe start/end positions placed the 50 µm nanoindentation grid points on the indented slice using gridline overlays to reconstruct the 50 µm-spaced nanoindentation grid on the indented section. These defined the measurement grid and yielded the ROI outlines in the coordinate frame of the indented section. Second, the cryosection IF and CytAssist images were aligned to the Visium HD fiducial markers in Loupe Browser, producing the JSON file used for bin alignment in Space Ranger. Third, the post-indentation confocal IF image of the indented surface was registered to the cryosection IF image by matching αSMA-positive vasculature features visible on both surfaces. Using the same antibody panel enabled direct comparison of these features. Translation, rotation, and scaling were determined independently for each tissue from the corresponding vascular. Fourth, the nanoindentation ROIs were transferred into the spatial transcriptomic coordinate system.

CytAssist-mediated sample transfer was restricted to tissue within the defined Visium HD capture area, so tissue extending beyond this boundary was not represented in the spatial transcriptomic dataset. Together with differences introduced by the adjacent tissue planes and their respective preparation workflows, this meant that the tissue areas represented in the nanoindentation images and final transcriptomic data did not overlap exactly. Accordingly, the complete nanoindentation ROIs shown in Extended Data Fig. 3 were not always represented in their entirety in the transcriptomic overlays shown in Extended Data Figs. 7 and 8; only grid points meeting the transcriptome-bin inclusion criterion were retrained for analysis.

Vascular-feature registration and transfer of the nanoindentation ROIs were performed in SpatialX v.2026-08-03 (BioTuring Inc), and each tissue-specific transformation was exported as a JSON file. This step can be performed in any spatial viewer with graphical alignment tools, such as napari (57) or STAlign (37). Operators were blinded to stiffness values and gene expression data, and transcriptomic cluster labels were not displayed during landmark selection. Each ROI was then delineated by point-to-point lasso selection tool, assigned a unique label in a dedicated ROI metadata column, and exported with the associated per-bin metadata as a tab-separated table.

### Assignment of transcriptome bins to nanoindentation grid points

Each indentation is indexed by tissue section, ROI, grid row, and grid column. A single 4 µm bin at each accessible ROI corner was annotated under a separate ROI-boundary metadata column using the convention <ROI-ID>_BotL/BotR/TopL/TopR. Two annotated corners sharing a grid row give the x-axis unit vector, scaled by the difference in their column indices, and two sharing a column given the y-axis vector. Both axes are forced orthogonal, and transcriptomic bin centroids were then projected into this coordinate system. Three annotations per ROI are sufficient to recover both axes, and all 21 ROIs met this. Recovery from incomplete corner sets is described in Supplementary Methods S8.

Each indentation was assigned the 4 µm bins projecting within its grid point, retaining up to 140 nearest the grid point centroid. Every bin assigned to an indentation inherits the same mechanical value, so constancy of that value across bins confirms the index is correct. Indentations retaining at least five bins were kept, giving 2,771 indentations across 8 ovaries and 21 ROIs, with recovered grid point dimensions of 30.4 to 51.0 µm and a median of 89 bins per indentation.

### Stiffness-expression association

Association between local mechanics and local transcription was assessed at the 50 µm nanoindentation grid point. Raw counts were summed across the 4 µm transcriptome bins assigned to each indentation and normalized once at that level, rather than correlating at 4 µm where most detected genes carry a single count (Supplementary Methods S9). Correlations were computed with Supplementary Code 12 and merged across analyses with Supplementary Code 13. Genes for spatial display were selected from the compartment differential expression tables, restricted to the Ovarian Mechanomics set, and plotted with Supplementary Code 14.

Associations are partial Spearman rank correlations between per-indentation aggregated gene expression and the mechanical parameter, holding sequencing depth and ovary identity constant. Expression, the mechanical parameter, and each covariate were rank-transformed, with the first two regressed on the covariate ranks by least squares and the residuals correlated. Depth is held constant because library-size normalization does not remove it from sparse data. Ovary identity is held constant because the 2,771 indentations derive from 8 ovaries and 6 animals.

Two age models were used. The pooled model controlled for age and depth, giving the stiffness association common to both groups (Fig. 3a). The stratified model computed the association within each age group separately, holding ovary identity constant within the stratum (Fig. 3d-i), and tested the difference between strata as an age-by-stiffness interaction by Fisher z transformation with the variance inflation appropriate to Spearman correlations. Ovaries are nested within age group, so age and ovary identity cannot both enter the pooled model. Both models were applied within each of the four compartments (Fig. 4e,h,k,n).

Every correlation was recomputed with each ovary removed in turn, and the range and smallest absolute value across folds are reported. Ranked mechanical variance is partitioned into between-ovary and within-ovary components for each parameter. Model detail is given in Supplementary Methods S10.

### Statistics and reproducibility

The experimental unit of observation depends on the question being tested and is stated in each case. The animal was the experimental unit for age-associated differential expression (*n* = 5 per group), the 50 µm nanoindentation grid point for mechanical association (*n* = 2,771 from 8 ovaries and 6 animals), and the individual indentation for descriptive mechanical statistics. Stiffness comparisons were made with linear mixed models on log_10_values, with animal, ovary, and region of interest as nested random effects (Supplementary Code 5). In the age-only model the animal-level variance component was negligible (0.009) against variance between ovaries (0.541) and between regions of interest (0.133), so the ovary is treated as the unit of replication. Compartment contrasts were Holm-corrected within three families, the pairwise compartment comparisons within young, those within aged, and the same-compartment comparisons between ages. Correlations at the measurement level are not independent within ovaries; this is addressed by covariate adjustment and by leave-one-ovary-out resampling rather than by treating measurements as replicates. Reported *P* values for partial correlations assume independent observations, which is not met, and are used for ranking rather than as calibrated error rates.

Age comparisons of mechanical parameters were made on the matched measurement set. ROIs were defined by overlap with the transcriptomic capture area and within the tissue interior to protect the probe, so they are not a random sample of the ovary, and the reported moduli describe the sampled regions rather than the organ as a whole. Random seeds are fixed throughout, and parallel and serial execution paths were verified to produce identical output. Registration and analysis controls are described in Supplementary Methods S11. Depth-confound diagnostics and registration controls were run with Supplementary Code 15 and 16.

## Acknowledgements

The authors thank the following people who contributed to this work: Arikta Biswas, Apoorva Shivankar, Boon Heng Ng, Haiyang Wang, Kim Whye Leong, Kosei Tomida and Lan Xi (MBI, NUS) for providing genes-of-interest in the ovarian mechanomics gene list; Anna Jaechke (MBI, NUS) and Apoorva Shivankar (MBI, NUS) for animal training and management support; Arikta Biswas, Kim Whye Leong, Anwesha Guru (MBI, NUS) and Avery Rui Sun (MBI, NUS; Stanford University) for fruitful discussions and technical advice; Shyam Prabhakar, Kok Hao Chen, and Nigel Chou (A*STAR) for discussions on the implementation of BANKSY and analysis approach; Anjali Verma and Quy Xiao Xuan Lin (10x Genomics) for Visium HD support; Raymond Woo (Carl Zeiss) for Axioscan 7 tissue scanner support; and Bournie Pham and Hazel Pham (BioTuring Inc) for SpatialX computational support. Special thanks to Raymond Rodgers (Adelaide University), Michael Stout (Oklahoma Medical Research Foundation), and Hattie Chung (Yale University) for their valuable time and expert insights on ovarian biology which improved this work. We thank the MBI Wet Lab Core for facility and technical support, the Singapore Microscopy and Bioimaging Analysis (SiMBA, MBI) core for microscopy and data processing facilities, and the High-throughput Molecular Genetics (HMG, MBI) core for molecular work support. We would also like to acknowledge Arikta Biswas, Kim Whye Leong, Anwesha Guru, and Raageshwari D/O Arulselvam (MBI, NUS) for reviewing and providing valuable feedback on the manuscript.

## Author contributions

Conceptualization: H.T.O and J.L.Y.

Experiments: H.T.O.

Analysis: Y.L. (Supplementary code 1), J.T. (Supplementary code 11 & 17), H.T.O. (all other scripts)

Contribution of unique tools or technical expertise: J.M. (visualization of methods into illustrations used in manuscript), R.M.H. (3D-printing and design of microscope slide chamber), M.F.H.R. (Support for tissue sample preparation for cryosectioning), X.S. (Support for tissue sample preparation for vibratome sectioning), J.Z. (Support for spatial transcriptome workflow and analysis), C.J.C. (Supplied animal models, vibratome usage, and support for tissue dissection)

Writing of manuscript: H.T.O., J.L.Y.

Revised manuscript: J.L.Y., C.J.C., R.L.

Supervision: J.L.Y.

Project support: C.J.C.

Funding and resources: J.L.Y., C.J.C., R.L.

Discussions: C.J.C., R.L.

## Declaration of interests and ethics

The authors declare no ethical or competing interests.

## Funding

This work was supported by the Bia-Echo Asian Centre for Reproductive Longevity and Equality (ACRLE) under the NUS Yong Loo Lin School of Medicine (J.L.Y., C.J.C., R.L.), National Research Foundation, Singapore, Mid-sized grant NRF-MSG-2023-0001 (J.L.Y., C.J.C., R.L.), Ministry of Education under the Research Centres of Excellence Programme through the Mechanobiology Institute at the National University of Singapore (NUS), the Biomedical Engineering Department at NUS (J.L.Y.), the Department of Biological Sciences at NUS (C.J.C.), Singapore Ministry of Education Academic Research Fund Tier 3 (MOET32021-0003 to J.L.Y, R.L.), Singapore Ministry of Education Academic Research Fund grant (MOE-T2EP30125-0011 to J.L.Y. and T2EP30222–0026 to C.J.C.), the support of the Singaporean Teaching and Academic Research Talent Inauguration Grant (to C.J.C.), Fudan University (Y.L.), Eric and Wendy Schmidt AI in Science Postdoctoral Fellowship (J.T.).

## Data availability

The data and images supporting the findings of this study are available within the article and its Supplementary Information. All raw and processed Visium HD spatial transcriptome data have been deposited as GEO accession GSE346700.

## Code availability

Analysis code is provided as Supplementary Code 1 to 18 (usage summarized in manifest README.txt), comprising nanoindentation force-curve processing and quality control (Supplementary Code 1 to 4), stiffness statistics (Supplementary Code 5), stiffness map generation (Supplementary Code 6), spatial transcriptomic bin size comparison (Supplementary Code 7), multi-sample integration and cell identity annotation (Supplementary Code 8), assignment of 4 µm transcriptome bins to 50 µm nanoindentation grid points (Supplementary Code 9), pseudobulk differential expression and gene set enrichment (Supplementary Code 10), curation of gene set enrichment plots (Supplementary Code 11), stiffness-expression correlation analysis (Supplementary Code 12), merging of correlation results (Supplementary Code 13), spatial plotting (Supplementary Code 14), the sequencing depth diagnostic (Supplementary Code 15), registration positive controls (Supplementary Code 16), til correction of post-nanoindentation confocal stacks in Fiji (Supplementary Code 18), and an extras folder (Supplementary Code 17) containing elastic net regression of stiffness on cell type composition, which was run on this dataset, explained almost none of the variation, and is provided for users to test on other tissue or disease models, and tilt correction of post-nanoindentation confocal stacks in Fiji (Supplementary Code 18). Random seeds are fixed throughout, and parallel and serial execution paths were verified to produce identical output.

The analysis code used in this study is available at https://github.com/huanting-ong/SpatialMechanomics, release v1.0.0, commit eb25d68, archived at Zenodo under DOI 10.5281/zenodo.22825608. The repository is released under an MIT license and includes Python and R environment specifications and a manifest README describing each script.

Supplementary Code 1: 1_NI_processing_pipeline.py

Supplementary Code 2: 2_NI_qc_indentation_depth.py

Supplementary Code 3: 3_NI_qc_sample_summary.py

Supplementary Code 4: 4_NI_parameter_summary.py

Supplementary Code 5: 5_NI_stiffness_stats.py

Supplementary Code 6: 6_NI_stiffness_map.py

Supplementary Code 7: 7_ST_Comparing2_4_8um.py

Supplementary Code 8: 8_Multisample-integration-VisiumHD.rmd

Supplementary Code 9: 9_Corr_roi_bin_assignment.py

Supplementary Code 10: 10_DEG_GSEA_YvA_wMechanics.Rmd

Supplementary Code 11: 11_curate_gsea_dotplots.py

Supplementary Code 12: 12_Corr_stiffness_expression_correlation.py

Supplementary Code 13: 13_merge_correlation_hits.py

Supplementary Code 14: 14_plot_segmentation.py

Supplementary Code 15: 15_depth_confound_diagnostic.py

Supplementary Code 16: 16_positive_controls.py

Supplementary Code 17: 17_Extra_celltype_mechanics_regression.py

Supplementary Code 18: 18_tilt_correction.ijm

## Extended Data Figures

**Extended Data Fig. 1.**
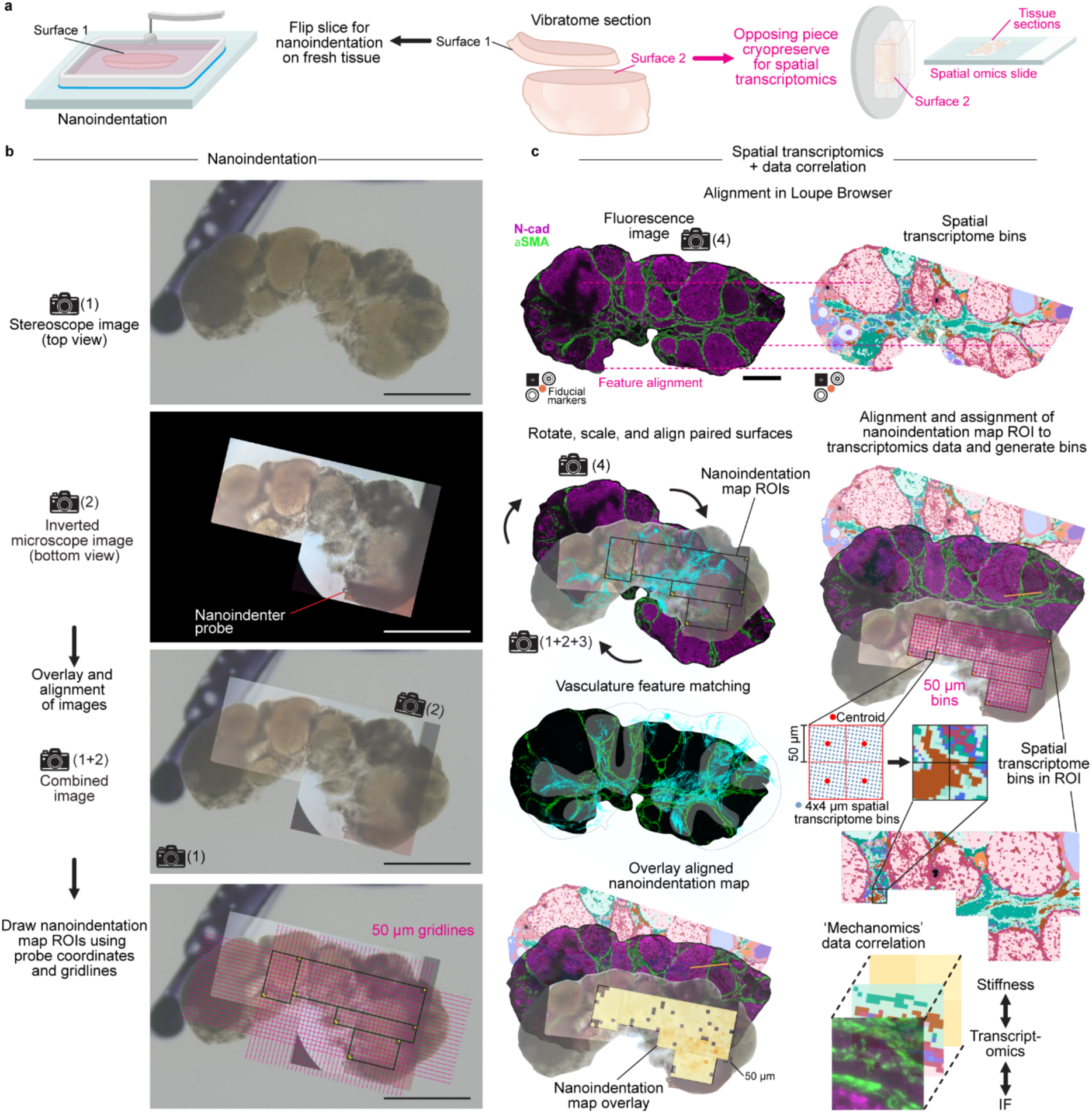
Worked example of paired-surface co-registration. (**a**) Opposing faces generated by a single vibratome cut were assigned to fresh tissue nanoindentation on surface 1 and cryosectioning for spatial transcriptomics on surface 2. (**b**) Top-view stereoscope (image 1) and bottom-view inverted microscope (image 2) images were aligned to locate the recorded probe positions and reconstruct the 50 µm nanoindentation grid within each ROI. Scale bar: 1 mm. (**c**) Cryosection IF (image 4; N-cadherin: magenta, αSMA: green) was aligned to Visium HD coordinates on Loupe Browser. The post-nanoindentation confocal IF image (image 3) was then rotated, scaled, and aligned to the cryosection IF image by matching αSMA-positive vasculature features. The nanoindenation ROIs were transferred into transcriptomic coordinates in SpatialX. The transformed grid, 4 µm spatial transcriptome bins, and Young;s modulus overlay are shown. Scale bar: 500 µm. This figure shows the complete registration workflow for one representative tissue.

**Extended Data Fig. 2.**
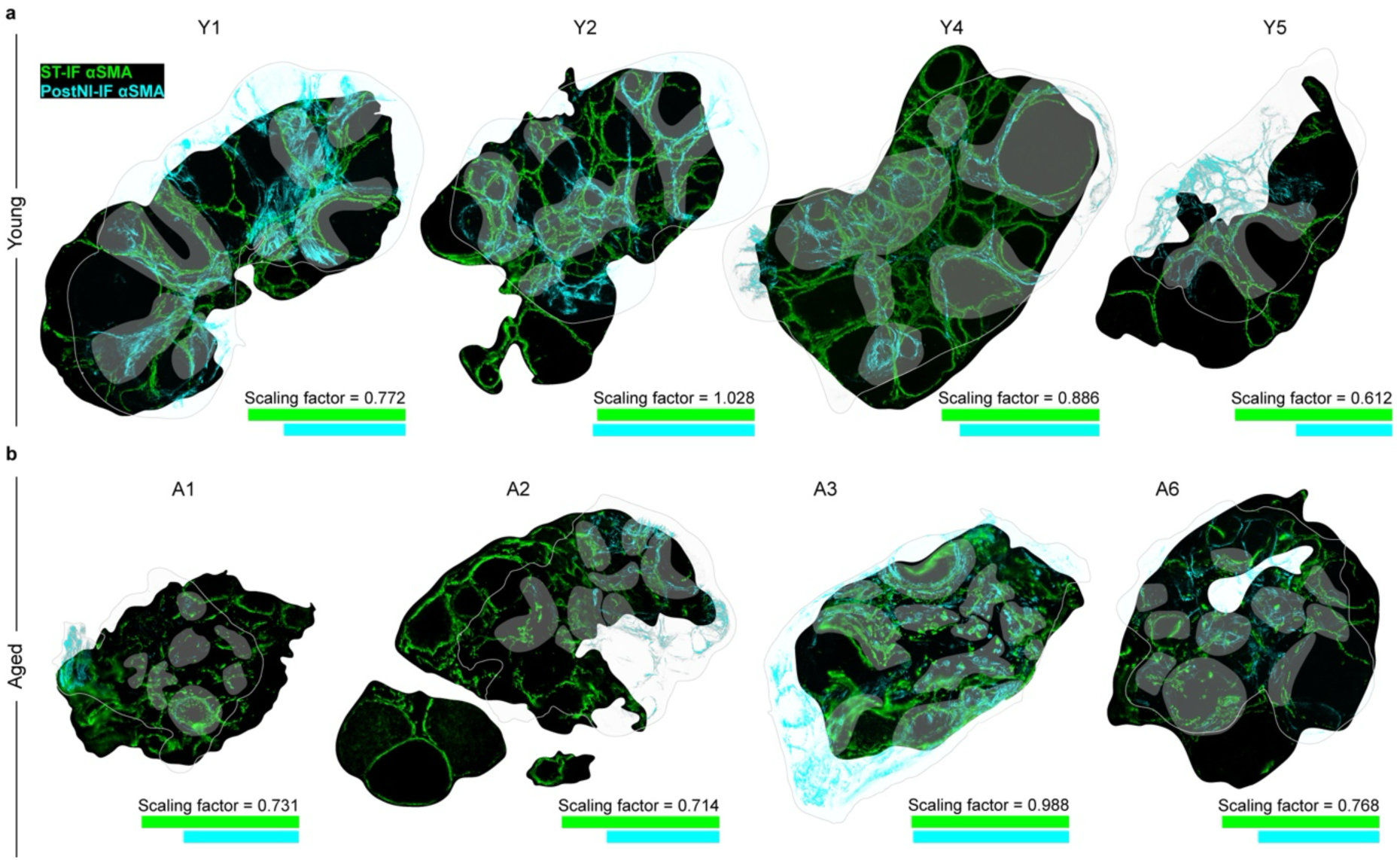
Vasculature features support registration of paired tissue surfaces across all eight ovaries. αSMA IF from the cryosection used for spatial transcriptomics (ST-IF, green) was overlaid with αSMA IF from the paired post-nanoindentation section (post-NI, cyan) after rotation and scaling for (a) four young and (b) four aged ovaries. The thin grey line marks the outline of the post-NI section. Overlapping green and cyan signals appear grey and identify vascular features represented on both surfaces. Because the images were obtained from adjacent tissue planes following different preparation and imaging procedures, exact correspondence of the tissue outline and smaller anatomical structures was not expected. Registration therefore used the distribution and branching patterns of conserved vascular features rather than complete tissue outline agreement. The dimensionless scaling factor obtained for each tissue is shown beneath its overlay and ranged from 0.612 to 1.028. Scaling was resolved independently for each tissue. Green and cyan scale bars: 1 mm.

**Extended Data Fig. 3.**
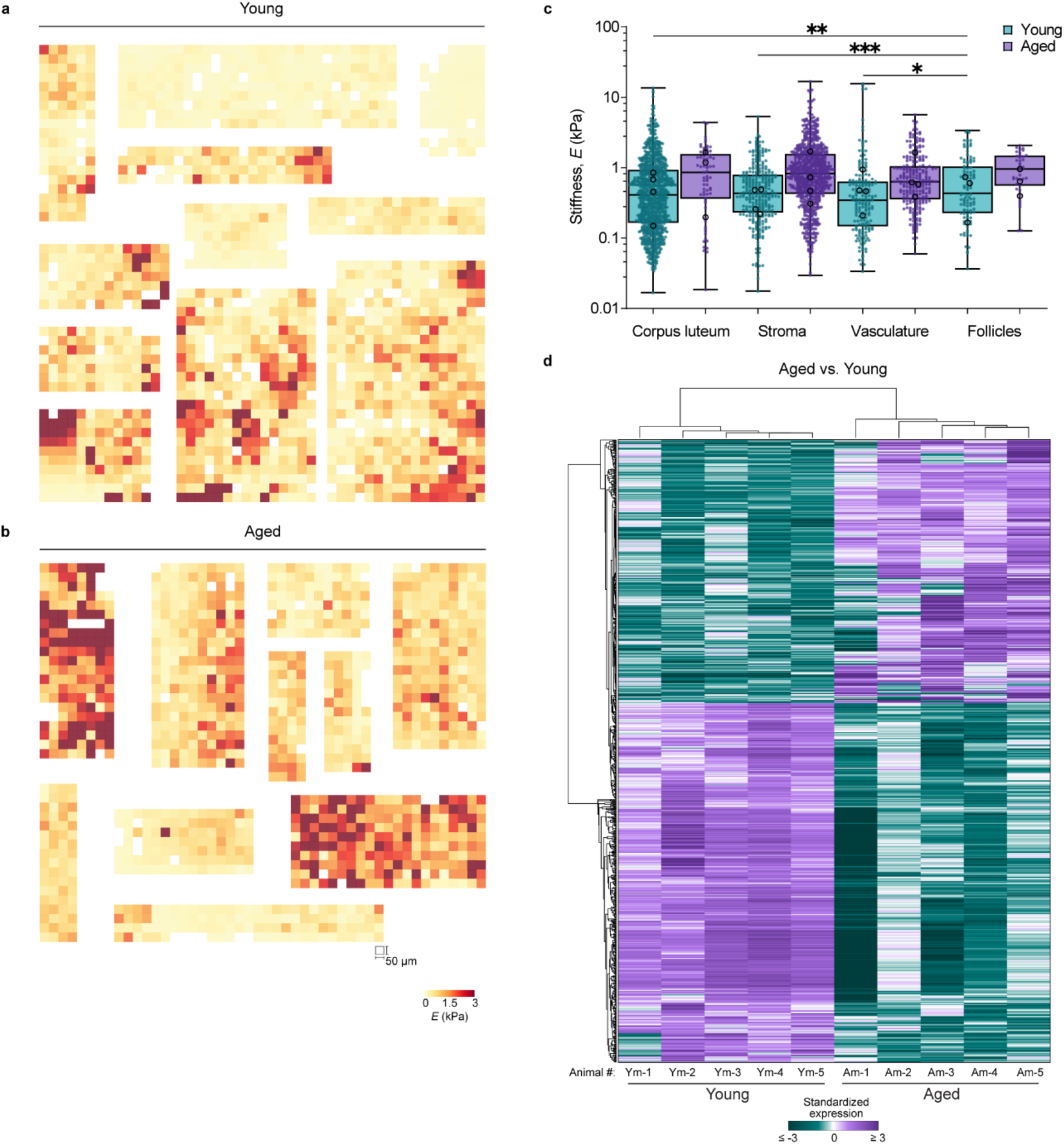
Stiffness maps, compartment stiffness comparisons, and age-associated gene expression across samples. (**a, b**) All matched ROI stiffness maps from young (**a**) and aged (**b**) ovaries. Each tile is one 50 µm nanoindentation grid point; white tiles denote measurements excluded during quality control. Young’s modulus scale: 0-3 kPa. (**c**) Stiffness separated by compartment and age. Box and whisker plots show median and interquartile range, with whiskers to min and max of all points within each compartment. Filled points represent individual grid points, and open circles represent per-ovary geometric means. Statistical comparisons used linear mixed model on log_10_-transofmred Young’s modulus, with animal, ovary, and ROIs included as nested random effects. Pairwise comparisons were Holm-corrected within the prespecified families. Brackets indicate comparisons within young tissue; no pair differed within aged tissue and no compartment differed between age groups (all Holm-corrected *P* > 0.6). The analysis included 1,648 young and 1,123 aged grid points from four ovaries per age group and six mice. \**P* < 0.05, \*\**P* < 0.01, \*\*\**P* < 0.001. (**d**) Heatmap of standardized expression for the 835 differentially expressed genes between aged and young tissue in the animal-level pseudobulk analysis. Columns represent 5 young and 5 aged animals, comprising 16 ovaries; rows represent genes. Rows and columns were ordered by hierarchical clustering using correlation distance. Differential expression thresholds were Benjamini-Hochberg *Padj* < 0.05, |log2FC| > 1.

**Extended Data Fig. 4.**
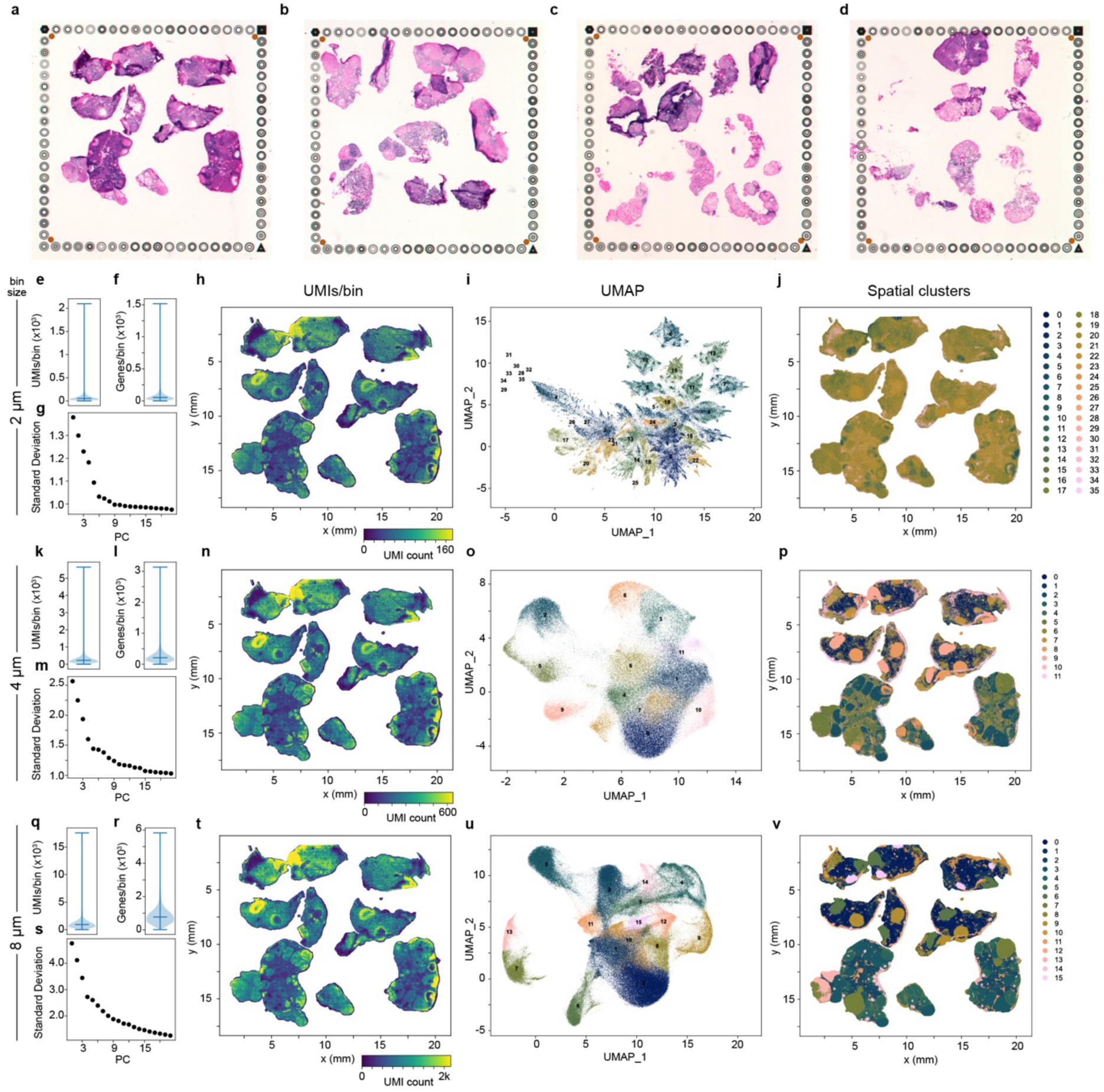
Visium HD quality control and bin size comparison. (**a-d**) H&E images of the four capture areas containing all 16 ovaries. (**e-j**) Results at 2 µm bin resolution: UMI counts per bin (**e**) number of detected genes per bin (**f**), variance explained by each principal component (**g**), spatial distribution of UMI counts per bin (**h**), UMAP representation (**i**), and spatial distribution of transcriptional clusters (**j**). (**k-p**) Corresponding results at 4 µm bin resolution. (**q-v**) Corresponding results at 8 µm bin resolution. The 4 µm bin resolution was selected for all downstream analyses as a balance between transcript capture and spatial detail.

**Extended Data Fig. 5.**
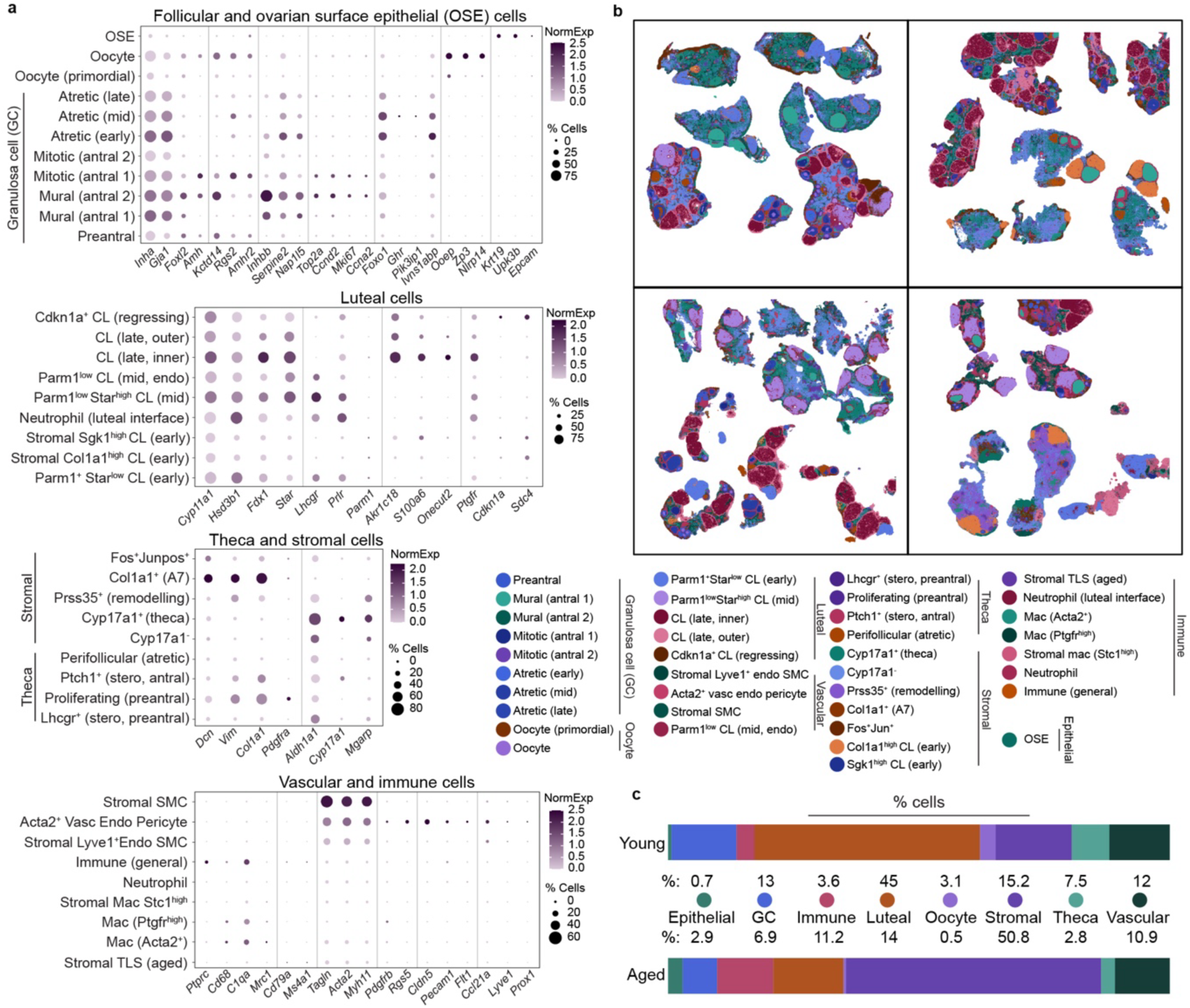
Marker gene and spatial evidence for cell identity assignment. (**a**) Dot plot of marker genes used to assign the 38 annotated cell populations. Dot size represents percentage of bins expressing each gene, and color represents scaled mean expression. Marker gene differential expression results are provided in Extended Data Table 2. (**b**) Spatial distribution of each annotated cell population, with the corresponding colors grouped by categories. (**c**) Percentage of bins assigned to each annotated population in young and aged tissue.

**Extended Data Fig. 6.**
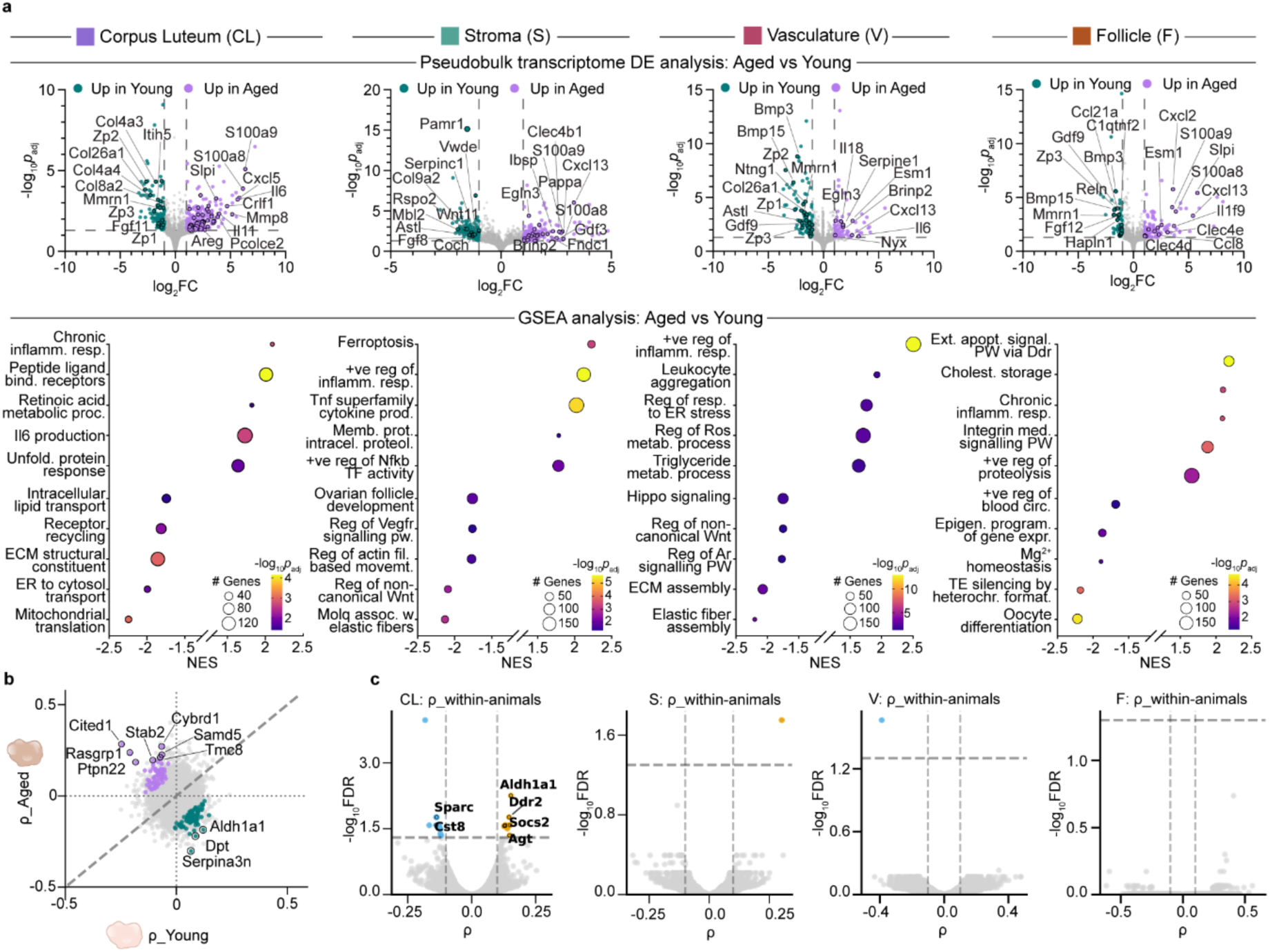
Compartment-specific age effects and local stiffness-expression associations. (**a**) Animal-level pseudobulk differential expression between aged and young tissue, computed separately within each compartment (corpus luteum, stroma, vasculature and follicle, from left to right). Counts were aggregated per animal, combining both ovaries where an animal contributed a pair; profiles containing fewer than ten bins were excluded. Dots colored in teal represent genes upregulated in young tissues and purple represent genes upregulated in aged. Gene set enrichment analysis was performed on the ranked differential expression statistic. Dot size represents the number of leading-edge genes, and dot color represents −log₁₀(adjusted *P*). Positive NES indicates enrichment in aged tissue. Gene sets were obtained from Hallmark, GO Biological Process, Cellular Component and Molecular Function, and Reactome collections; redundant sets were collapsed by leading-edge overlap and representative terms selected for display. (**b**) Partial Spearman correlation (ρ) in young tissue plotted against ρ in aged tissue, with one point per gene. The 146 genes meeting the prespecified threshold for a difference between age groups by Fisher *z* transformation are highlighted; 81 were classified as more strongly correlated in young tissue and 65 in aged tissue. The dashed diagonal marks denote equal ρ between both age groups, and dotted lines denote ρ = 0. (**c**) Compartment-resolved partial Spearman correlations between aggregated gene expression and Young’s modulus, with sequencing depth and ovary identity held constant. Correlations are plotted as −log₁₀(FDR) against ρ. Dashed lines denote FDR = 0.05 and |ρ| = 0.1. Labelled genes are those belonging to the curated ovarian mechanomics gene set; other genes meeting the thresholds are unlabeled for clarity.

**Extended Data Fig. 7.**
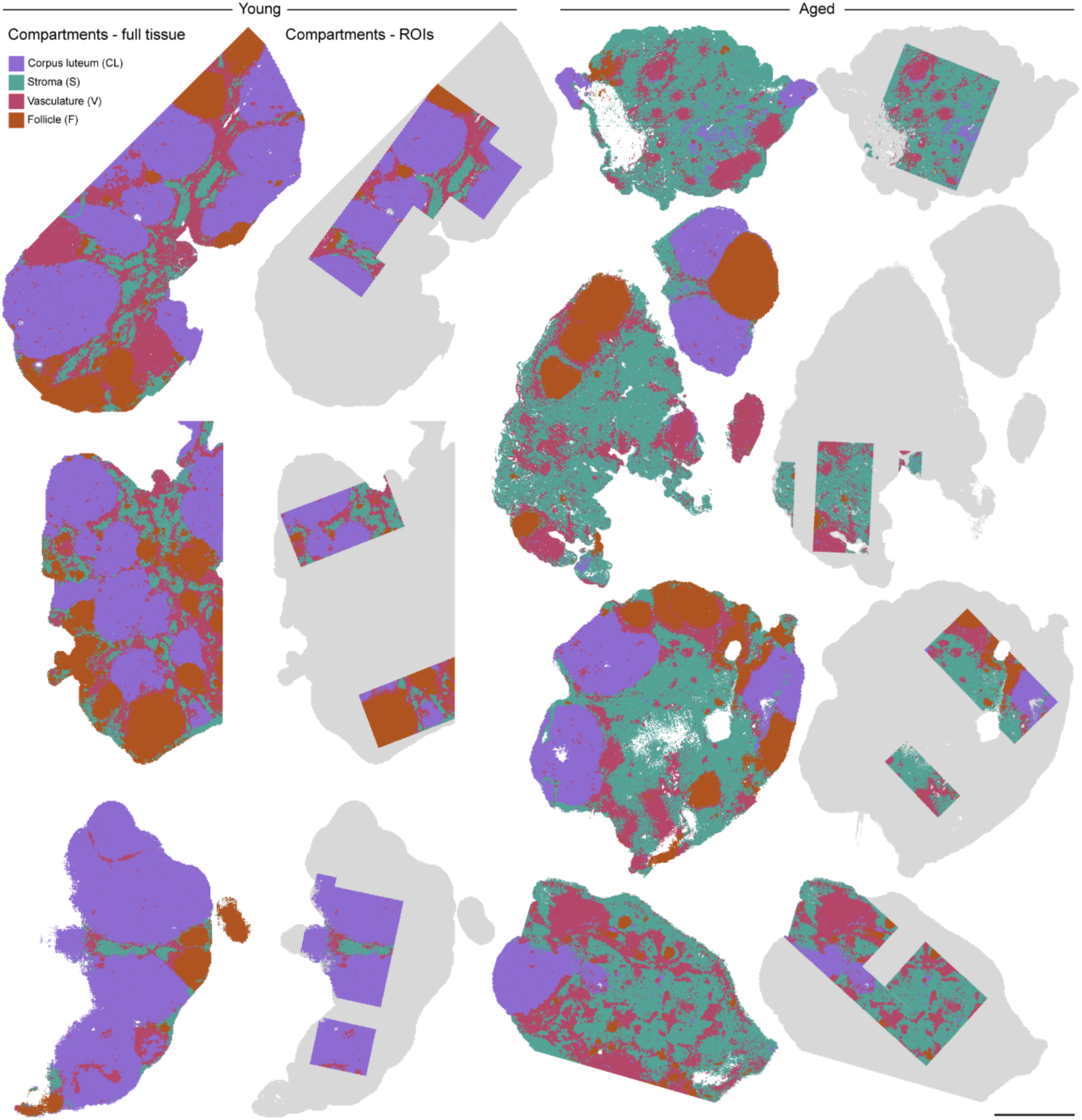
Compartment annotation of matched ROIs for all remaining ovaries. Full-section compartment maps and retained ROI masks are shown for young (**a**) and aged (**b**) ovaries not displayed in Fig. 4a. The tissue sections were annotated into the four major compartments: corpus luteum (CL) in purple, stroma (S) in teal, vasculature (V) in pink, and follicle (F) in brown. Tissue outside matched ROIs in grey, and regions without tissue regions are shown in white. Scale bar: 500 µm.

**Extended Data Fig. 8.**
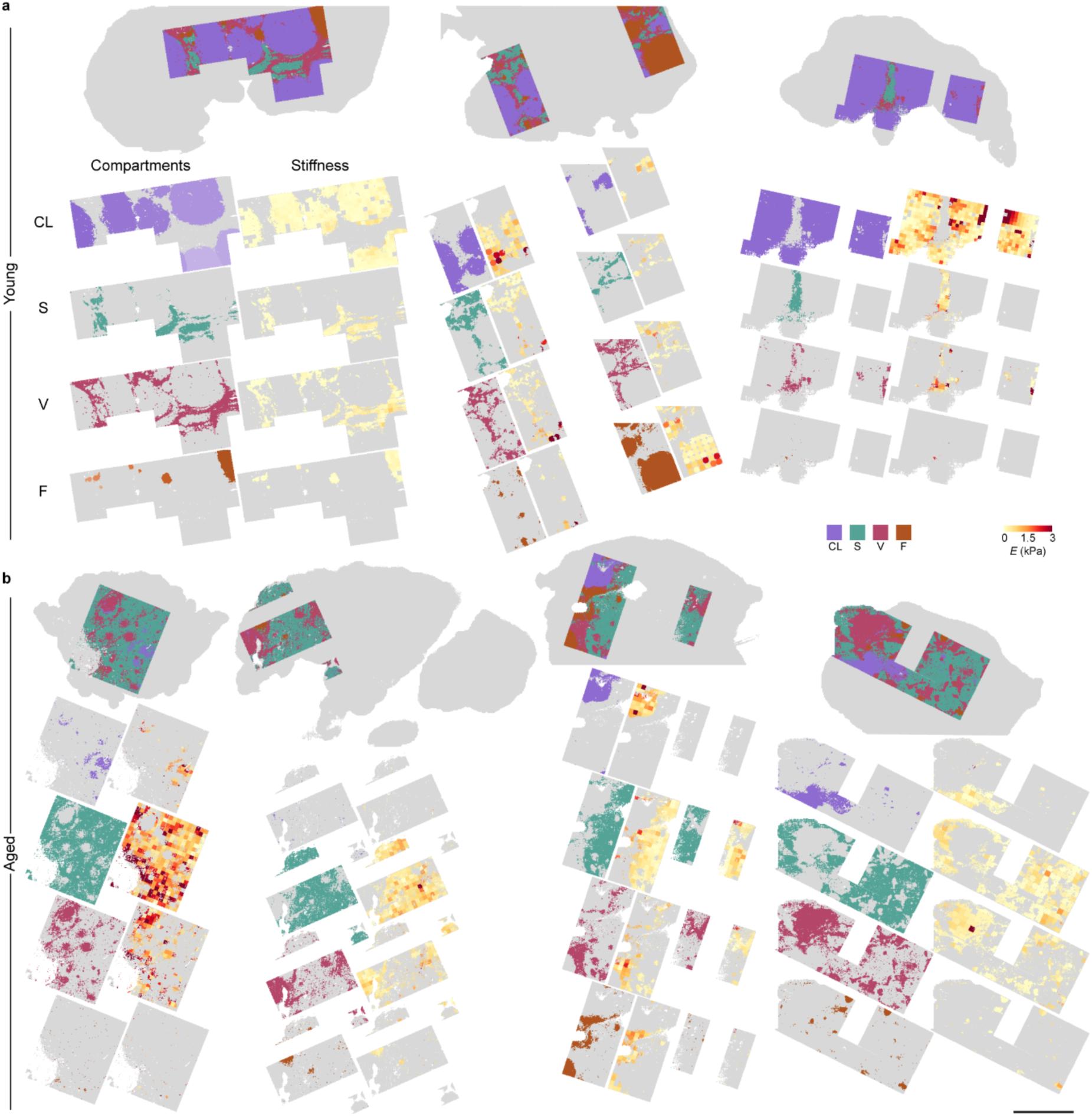
Compartment-resolved stiffness maps for all remaining ROIs. Compartment assignments and matched stiffness maps are shown for ROIs from the remaining young (**a**) and aged **(b**) ovaries displayed in Extended Data Fig. 7. For each tissue the upper map shows the compartment assignment with the complete matched ROI. The rows below show the bins assigned to corpus luteum (CL) in purple, stroma (S) in teal, vasculature (V) in pink, follicle (F) in brown,, together with the corresponding Young’s modulus values at the matched nanoindentation grid points. Tissue outside the matched ROIs is shown in grey, and regions without tissue regions are shown in white. Nanoindentation grid spacing: 50 µm; Young’s modulus scale: 0-3 kPa. Scale bar: 500 µm.

## Supplementary information

### Supplementary Methods

#### S1. Slide preparation, tissue attachment, indentation chamber, and cryosectioning

Microscope slides (#406/0179/00, VWR) were cleaned in isopropanol, rinsed in sterile-filtered ultrapure water, dried, and treated in a UV ozone cleaner (UV/Ozone ProCleaner Plus, BioForce Nanosciences) for 5-10 min to increase surface hydrophilicity. In a biological safety cabinet, 20 µl of sterile poly-D-lysine (PDL, 0.1 mg/ml, molecular weight 50,000 to 150,000 Da; #A3890401, Gibco) was applied to the center of each slide and allowed to dry completely, repeated two more times. Coated slides were covered and sealed at room temperature and used within 2 days or stored up to 1 week to exclude dust until use.

Before use, coated slides were rinsed three times in sterile-filtered ultrapure water and once in 1X PBS, and the coated region marked on the underside with a lab marker. Fresh vibratome sections of ovarian tissue were transferred with fine forceps via a clean transfer slide under a stereomicroscope, floated into position within a 1X PBS droplet over the marked region, and settled onto the coating with fine forceps.

Once attached, the slide periphery was dried while the tissue remained immersed in L-15 medium. A 3D-printed chamber wall of internal dimensions (16mm width × 39mm length x 5mm height) was fixed in position with duplicating silicone eco-sil speed (#13007100, Picodent Twinsil) mixed 1:1 and cured for 5 min at room temperature, and the chamber filled with L-15 medium. The 3D-printed-chambers were designed in Onshape (Onshape Inc., USA) and exported as STL files. Print files were generated using PrusaSlicer and fabricated using an Original Prusa SL1S SPEED 3D printer (Prusa Research, Czech Republic) at 25-μm layer resolution. Components were printed using Prusament Resin Model Alabaster White (Prusa Research, Czech Republic). Following printing, the chambers were washed in isopropyl alcohol to remove uncured resin before drying and UV post-curing with the Original UV according to the manufacturer’s recommendations. To further remove residual resin, 3D-printed chambers were sequentially washed in acetone (Sigma-Aldrich), ethanol (Sigma-Aldrich), and deionized water (Milli-Q, Merck Millipore). Before use, the pieces were sterilized in 70% ethanol and washed in deionized water and 1X PBS.

Tissue attachment was verified by topping up L-15 medium over the section: a section that remained in place rather than lifting under its own buoyancy was taken as attached. Sections that failed to attach were discarded and a further section taken from the same ovary onto a fresh coated slide, since the coating can be lifted away with the discarded tissue. The 1 mm parallel block was cut only after a section had been confirmed attached, so the cryopreserved face is in every case the face immediately beneath the section that was indented.

For cryosectioning, in order to ensure the block face is parallel to the cutting plane prior to reaching the tissue block, an alignment step was carried out. On top of the OCT block containing the tissue, a droplet of OCT was applied, which freezes into a dome. If the blade is not parallel to the mounted face, the dome will produce an elongated ellipse section when cut rather than a circle. The shape of the OCT profile during trimming was therefore used as a running check on alignment, and trimming was adjusted until sections came off circular. Every section once the tissue became visible underneath the OCT was examined under the microscope during trimming, to ensure the tissue surface 2 would be captured. Once surface 2 was reached, the first intact 10 µm section was used for Visium HD analysis; subsequent serial sections were collected as backups in case of section damage.. The serial sections were placed onto a pre-marked Superfrost^®^ Plus Adhesion microscope slide (#406/0179/00, VWR) to fit within a 6.5 mm × 6.5 mm marked region corresponding to the Visium HD capture area and processed using CytAssist according to the manufacturer’s workflow, summarized in Extended Data Table 7.

### S2. Derivation of viscoelastic parameters

The pipeline additionally returns the effective modulus fitted to the collapsed loading and retraction curves, the modulus fitted to the retraction curve alone, viscosity under a Maxwellian model, a loss ratio giving the fraction of energy dissipated per load-unload cycle, and the loading, retraction and adhesion energies. These are written to the output tables and available for reuse, but are not analyzed here. The implementation follows Biswas et al. (21). Under the Maxwellian model, the force *F* at the tip is assumed to balance by the elastic component and viscous component in series, so the displacement δ is the sum of the elastic and viscous contributions. Hence, the force measured by the tip *F* and tip displacement δ during loading and retraction in the positive force region are assumed to obey the following evolution over time *t*, respectively:

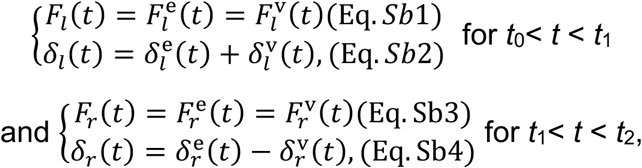

with boundary conditions *F_l_*(*t*_0_) = *F_r_*(*t*_2_) = 0, *F_a_*(*t*_1_) = *F_r_*(*t*_1_), *δ_a_*(*t*_0_) = 0, *δ_a_*(*t*_1_) = *δ_r_*(*t*_1_). All the superscript *e* and *v* denote the elastic or viscous counterparts. For simplicity, we assume a linear relationship between viscous force and viscous displacement 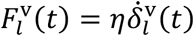 and 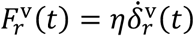, with the viscosity coefficient *η* in the unit of Ns/m, and the elastic contribution follows the same Hertzian relationship between elastic force and elastic displacement 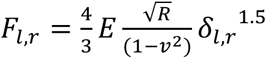, where *E* is the elasticity of the elastic component.

Substituting Eq. Sb1(or Sb3) into Eq. Sb2(or Sb4) gives us the elastic component displacement which can be related to force as

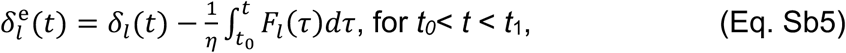

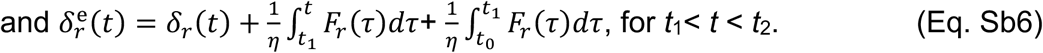

The viscous force, 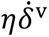, is assumed as linear to the shrinkage rate of the viscous damper 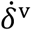, to be measured from the data. The displacement of the viscous components is calculated as 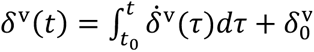, where 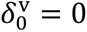 for *t*_0_. Note that *F*(*t*_2_) =0, the elastic component at *t*_2_ also has zero displacement. From Eqs. Sb1-2, the distance between the tip position *δ_r_*(*t*_2_)-*δ_l_*(*t*_0_) is derived to be

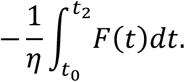

From the data, we could directly measure *δ*_r_(*t*_2_) and *δ*_l_(*t*_0_) and thus obtain the Maxwellian viscosity of the system.

Two further quantities are derived from the same curves without reference to the Maxwellian model. The energy dissipated through a whole process of loading and retraction is calculated as the difference between the loading work *W_l_* and the work released by retraction *W_r_*, where the work *W_l,r_* is *F_l,r_* integrated over loading/retraction displacement for the range of *F_l,r_* >0 . Then, the loss ratio, *H* is defined (*W_l_ -W_r_*)/*W_l_*, indicating the extent of non-elasticity (viscosity and plasticity) of the tissues. Adhesion energy was obtained by the calculating the work by negative retraction force, which is the area enclosed by retraction curve for *F_r_* <0. This represents adhesive interactions between the probe and the tissue surface, likely influenced by ECM proteins and hydration.

#### S3. Nanoindentation curve quality control (QC)

The following criteria were applied before a curve contributed a mechanical value, two acting on the raw force-displacement data, one on the Hertzian fit, and one on the fitted modulus. Two further constraints were applied to derived parameters.

##### Failed indentations

Files reporting a maximum load of −0.001 µN were treated as no indentation and excluded.

##### Contact detection

Two models were fitted to each approach segment: a single straight line, and a piecewise model of a flat pre-contact baseline followed by a Hertzian indentation regime. The RSS ratio is the residual sum of squares of the former divided by that of the latter, and curves were retained above a threshold of 1.655. On 39 manually labeled curves from six tissue types the two classes separated completely, with no-contact curves spanning 0.182 to 1.082 and contact curves 2.530 to 394; 1.655 was taken as the midpoint of the gap. The two criteria agreed on all 1,870 points across the full dataset where the instrument reported its own maximum-force diagnostic.

##### Hertzian fit quality

Curves were retained at a relative amplitude standard error (Perr_l) ≤ 0.15. Relaxing this from the conventional 0.05 changed the pooled median modulus by 1.6%, recovered 4% additional coverage, and reduced the differential loss between age groups from 0.84 to 0.26 percentage points. The aged-to-young median ratio was 1.602 at 0.05 and 1.647 at 0.15.

##### Modulus bounds

Fitted moduli outside 7.6 to 22,726 Pa were excluded, these being the limits of the measurable range for this probe as determined from its calibrated spring constant and tip radius.

##### Viscosity

Raw viscosity spanned many orders of magnitude with no corresponding signature in any fit-quality metric, so a magnitude cap of 1.0 N s/m was applied. Neither the amplitude standard error of the loading fit nor that of the collapsed fit discriminates implausible viscosity values, since the latter quantifies agreement between loading and retraction curves under a single Hertzian model rather than the fidelity of a viscoelastic fit.

##### Asymmetry between elasticity parameters

*E* is masked by Perr_l. *E*_eff_ has no equivalent mask, because the corresponding error term is not a fit-quality criterion for the effective modulus. Comparisons involving *E*_eff_ are therefore made under weaker quality control, which is one reason *E* is the primary parameter throughout.

#### S4. Bin size selection

Visium HD data can be aggregated to any bin size that is a multiple of the 2 µm capture square, and the choice sets the effective resolution of every downstream analysis. Three candidate resolutions (2, 4, and 8 µm) were compared directly on these data before any biological analysis was run (Extended Data Fig. 4, Supplementary Code 7).

Each capture area was processed independently at 2, 4 and 8 µm, comparing per-bin UMI counts, per-bin detected features, principal component structure, cluster separability in UMAP and the spatial coherence of clusters. Bin counts per capture area were 4,388,864, 4,145,192, 4,169,800 and 3,028,592 at 2 µm; 1,097,216, 1,036,298, 1,042,450 and 757,148 at 4 µm; and 274,304, 259,204, 260,642 and 189,322 at 8 µm, over 19,059 genes (Extended Data Table 1). At 2 and 4 µm the bin count exceeds what can be clustered directly, so clustering was performed on a 150,000-bin subsample and labels projected back to all bins by nearest-neighbor matching in principal component space, allowing spatial coherence to be assessed on the full tissue.

Cluster number at a fixed resolution of 0.6 was not monotonic in bin size: 30 to 49 clusters at 2 µm, 10 to 12 at 4 µm and 15 to 17 at 8 µm. The excess at 2 µm reflects sparsity rather than biology, since most 2 µm bins carry only a handful of UMIs and split on noise. The apparent deficit at 4 µm relative to 8 µm is an artifact of the subsampling and projection step used at the two finer resolutions and does not indicate loss of information; the 38 populations reported in this study were resolved from the integrated four-capture-area object at resolution 0.8, not from these per-capture-area comparisons, and those two cluster counts are not comparable.

4 µm was selected because: 1) it retains enough counts per bin for stable clustering, which 2 µm does not, and 2) it resolves features that 8 µm merges: the neutrophil population at the early corpus luteum border occupies two to three contiguous 4 µm bins, which fall within a single bin at 8 µm.

#### S5. Bin population cell annotation strategy

Populations were annotated using a multi-evidence framework applied to the 38 clusters retained after removal of five clusters comprising fewer than 50 bins, which were excluded because they could not be annotated robustly. For each cluster a marker gene score was calculated as MG_score = avg_log_2_FC × (pct.1 − pct.2) × (−log_10_(*P*_adj_ + 1 × 10⁻³⁰⁰)) from one-versus-all Wilcoxon rank-sum testing (Extended Data Table 2). Where one-versus-all comparisons could not resolve closely related populations, pairwise differential expression was performed in SpatialX with the same test at |log_2_FC| > 1. Spatial expression patterns were interpreted against known tissue architecture, including follicle morphology, corpus luteum stage and stromal organization. Co-localization with αSMA immunofluorescence and age-specific distribution across capture areas provided additional evidence, and cluster stability was assessed by clustree across resolutions.

##### Follicle compartment

Two oocyte populations were distinguished by stage. Primordial oocytes (cluster 22) express the oocyte fate transcription factors *Sohlh1*, *Sohlh2*, *Figla*, *Nobox*, and *Lhx8*, with the meiotic marker *Sycp3*, and appeared as discrete cortical foci. Growing preantral and antral oocytes (cluster 26) express *H1foo*, *Gdf9*, *Zp3*, *Nlrp14*, and *Mos*, and localize to follicle centers. Granulosa cell populations share *Inha*, *Gja1*, *Foxl2*, and *Amh*, and subdivide by stage: preantral (cluster 14, *Kctd14*, *Rgs2*, and *Amhr2*), a spatially intermediate cumulus population (cluster 21) positioned between the preantral oocyte cluster and the outer granulosa layer, mitotic preantral (cluster 23; *Top2a*, *Ccnd2*, *Mki67*, *Ccna2*), antral mural (clusters 11 and 34; *Inhbb*, *Serpine2*, *Nap1l5*) and mitotic antral (cluster 30). Atretic granulosa populations (clusters 28, 19 and 32) were staged early, mid and late on progressive *Foxo1*-mediated reprogramming, marked by *Foxo1*, *Ghr*, *Pik3ip1* and *Ivns1abp* rather than by generic apoptosis signatures. Cluster 18 occupies a perifollicular position around atretic follicles but was separated from atretic granulosa by pairwise testing showing upregulation of *Bgn*, *Dcn*, *Aldh1a1*, *Cyp11a1* and *Prss35* and absence of *Fst* and *Nap1l5*, confirming a stromal-theca identity.

##### Corpus luteum compartment

Early corpus luteum comprises cluster 7 (*Parm1*-high, *Star*-low, indicating luteinization before full steroidogenic activation) and two stromal populations separated by pairwise testing, cluster 13 (*Col1a1*-high, ECM scaffold) and cluster 37 (*Sgk1*, stress or hormone-responsive). Mid corpus luteum comprises cluster 5 (*Parm1*-low, *Star*-high, peak steroidogenesis) and cluster 15, a luteal endothelial population. The late corpus luteum inner core (cluster 1) expresses *Hmgcr* and *Scd1*, consistent with maximally steroidogenic granulosa-lutein cells, and the late outer border (cluster 3) expresses *Aldh1a1*, *Acta2* and *Dcn*, consistent with theca-lutein cells. Two macrophage populations occur in the late corpus luteum, cluster 9 (*Acta2*-positive) and cluster 25 (*Ptgfr*-high, consistent with a luteolysis-associated state). Regressing corpus luteum (cluster 27) co-expresses *Cdkn1a*, *Spp1*, *Wnt10b* and *Cemip*, consistent with p21-mediated arrest and active ECM remodeling.

##### Stromal compartment

Clusters 0 and 2 were separated by pairwise testing showing downregulation of *Cyp17a1*, *Fabp3*, *Adh7*, *Fads2* and *Agt* in cluster 0; cluster 2 was annotated as a steroidogenically active theca-interstitial population dominating aged stroma, and cluster 0 as a quiescent stromal equivalent present at both ages. Clusters 8 and 35 share *Ptch1* and *Inhba*, consistent with hedgehog signaling from adjacent granulosa cells, and were distinguished by association with antral and preantral follicles respectively. Cluster 12 was annotated as an active ECM-remodeling stromal population on *Prss35*, *Adamts1*, *Pcsk5*, *Timp1* and *Plau*. Cluster 36 (*Col1a1*-high) was restricted entirely to one tissue slice and is retained with that restriction flagged.

##### Vascular and smooth muscle compartment

Cluster 4 expresses *Ccl21a*, *Acta2*, *Myh11* and *Col1a1* and is interpreted as a composite vascular-stromal niche reflecting the spatial co-organization of lymphatic endothelium, smooth muscle and ECM-rich stroma at 4 µm rather than a transcriptionally pure cell type. Cluster 10 co-expresses the endothelial markers *Cldn5* and *Epas1*, the pericyte marker *Rgs5*, and *Ccl21a* with *Tm4sf1*, reflecting interstitial vasculature with mixed endothelial and mural signal. Cluster 20 is defined by *Actg2*, *Lmod1*, *Cnn1*, *Des* and *Tagln*; the specificity of *Actg2* for visceral rather than vascular smooth muscle distinguishes it from cluster 10. Both are αSMA-positive by IF.

##### Immune compartment

Cluster 17 occurs exclusively in aged stroma as discrete focal aggregates, spatially consistent with tertiary lymphoid structures. Its dominant signature of immunoglobulin genes (*Igkc*, *Igha*, *Ighm*, *Ighg1*, *Ighg2b*, *Jchain*) identifies the constituent cells as plasma cell-rich aggreagates spatially consistent with TLS-like structures, consistent with the age-related expansion of ovarian lymphoid populations reported in the mouse ovarian single-cell atlas (23) and with tertiary lymphoid structure accumulation in aging tissues generally (33,34). Clusters 29 and 33 colocalize exclusively at early corpus luteum structures in a recurring pattern, with single cluster 29 bins encircled by cluster 33 bins. Cluster 29 carries a canonical neutrophil signature (*S100a8*, *S100a9*, *Camp*, *Ltf*, *Ngp*, *Lcn2*, *Csf3r*), consistent with polymorphonuclear neutrophil infiltration into the developing corpus luteum (32), and cluster 33 captures the mixed neutrophil-luteal bins localized to the interface. Given a mouse neutrophil diameter of 10.1 to 10.6 µm (57), this interaction zone spans two to three contiguous 4 µm bins and would be masked within a single 8 µm bin. Cluster 24 (*Stc1*-high) was restricted to one capture area and is flagged. Cluster 31 comprises residual mixed immune populations not meeting the threshold for discrete annotation.

##### Flagged annotations

Cluster 6 expresses the immediate early genes *Fos* and *Jun* at high levels across both percentage of expressing bins and log₂FC relative to all other clusters. This pattern is a documented artifact of tissue stress, but was retained given the possibility of biologically meaningful immediate early gene expression in aged tissue under physiological stress.

#### S6. Pseudobulk differential expression and curated gene sets

Profiles are formed by summing raw counts, and the number of 4 µm bins contributing to a profile ranges over more than two orders of magnitude between ovaries for some cell clusters. A profile from few bins is noisier than one from many, which affects dispersion estimation in DESeq2. The minimum-bin threshold used is stated in the Online Methods. Two adjusted P values were computed and retained, one from the ordinary Wald test used throughout and one from a threshold-based test kept for reference. The threshold-based version returns adjusted P equal to one for a large block of genes and cannot be used for ranking. The differential expression analysis uses all 16 ovaries. The mechanical analyses use the 8 for which nanoindentation was performed.

Differentially expressed genes were classified against the mouse matrisome masterlist (35), which resolves into two divisions (core matrisome, matrisome-associated) and six categories (collagens, ECM glycoproteins, ECM regulators, ECM-affiliated proteins, proteoglycans, secreted factors). An “Ovarian Mechanomics” gene set of 1,854 symbols was additionally assembled by merging the matrisome masterlist with a curated list of mechanobiology and ovary-specific genes, including granulosa apoptosis and follicle growth genes, and adding binding partners recovered from a ligand-receptor pair database for genes not already on either list. Ligand-receptor entries with no matrisome or curated-list partner were excluded. Of the 1,854 genes, 1,724 were present in the expression matrix. Full membership and provenance per gene are given in Extended Data Tables 5 and 6.

#### S7. Gene set enrichment and pathway redundancy

The ranking statistic for enrichment on differential expression is the signed Wald statistic from the unshrunken model, which is continuous across the full gene list and preserves both direction and magnitude of evidence. Permutation-based significance was estimated with a fixed random seed. Gene set size limits define which sets are tested and therefore also the size of the multiple-testing family, so adjusted *P* values depend on the size window for every pathway and not only for those added or removed. The window was fixed before results were inspected.

Leading-edge clustering groups significant pathways whose leading-edge gene sets overlap above a stated Jaccard threshold, reports one representative per cluster together with the genes shared by at least half its members, and records the full membership. This is necessary because MSigDB set names reflect the experimental context in which genes were first characterized rather than the tissue under study, so pleiotropic signaling and immune genes generate enrichment of terms with no ovarian interpretation. Where the same leading edge recurs across compartments, the finding is reported once as a shared program rather than as independent enrichment in each compartment, since the compartment tests share ovaries and the same infiltrating populations contribute bins to more than one compartment.

Enrichment on the stiffness-expression correlations was computed by preranked GSEA on the full ranked list of ρ values. Only gene sets with 10 to 500 members present in the ranked list were tested. The gene set libraries used for this analysis were GO Biological Process and Reactome terms.

#### S8. Bin assignment implementation

Where fewer than three annotations exist, principal component analysis of the ROI bin cloud supplies the axes, mapping the longer principal component to the longer grid dimension, with the single annotation fixing the sign. This is unreliable for grids close to square and the pipeline reports when it is used. No ROI in this study required it. Grid-cell boundaries were defined from the annotated ROI corners and the recovered row and column vectors; the observed transcriptomic bin cloud was then projected into these fixed boundaries. The composite index is necessary because grid coordinates repeat across ROIs and slices. Grouping on the coordinate alone collapsed 2,789 distinct indentations into 521 groups, of which 82% contained more than one distinct modulus value. ROI labels follow the convention <OVARY=-<INDEX=, for example Y4-2. The pipeline runs on Python 3.13.13 with NumPy, Pandas and Matplotlib, and exports the enriched metadata table as a TSV.

#### S9. Sequencing depth confounds bin-level correlation

Bin-level correlation between gene expression and stiffness was computed and rejected. Log-normalized expression is log(1 + c/N × s), where c is the raw count for a gene in a bin, N the bin’s total count and s a global scale factor. At 4 µm most detected genes have c of 1 or 2, so conditioning on detection leaves variation dominated by 1/N, which is shared across genes. Under that condition ρ(expression, E) approximates −ρ(N, E) for almost every gene.

The number of genes detected per bin correlated with log-transformed Young’s modulus at ρ = −0.248 across all 255,620 bins carrying a valid modulus, and −0.185 to −0.233 across compartments. Per-gene correlations were computed three ways on the same 1,048 genes: conditioned on detection, 99.6% were positive, with a median of +0.246; on all bins with zeros included, 1.2% were positive with a median of -0.037; with the number of detected genes partialled out of both variables, 90.1% remained positive but with a median of +0.028. The sign of the result therefore depends on how detection is handled, and the magnitude collapses when this single technical covariate is removed.

The same mechanism explains a related summary statistic: mean expression across the genes detected in a bin correlated with the number of detected genes at ρ = −0.9994, because library-size normalization fixes the un-logged total per bin. That quantity measures library complexity, not expression level.

Aggregating bins to the 50 µm nanoindentation bin raises the typical per-gene count out of the single-count regime and is the basis of the analysis reported. At point level the same 1,048 genes gave 55.5% positive correlations with a median of +0.009, and 536 reached FDR < 0.01 against 1,047 at bin level. It also raises the number of testable genes: 1,048 genes passed the bin-level detection filter of 5% of bins, against 11,255 passing the point-level filter of detection in at least 10% of points with at least 20 summed counts. Diagnostic code is provided as Supplementary Code 15.

#### S10. Point-level correlation model

Counts were summed across the 4 µm-equivalent stiffness bins of each nanoindentation and normalized once at that level. Expression is normalized within each bin by dividing a gene’s raw count by the bin’s total count, then log-transforming. At 4 µm most detected genes have a raw count of one, so the normalized value is set almost entirely by the denominator, which is shared across genes in that bin. Summing counts across the bins raises the per-gene count out of that regime. Where a raw-count export was unavailable, counts were recovered exactly by inverting the log-normalization using the per-bin total count. Recovered values are integer multiples of a single global constant, which was verified before use and cancels on renormalization.

Partial Spearman correlation was computed by rank-transforming expression, the mechanical parameter and each covariate, regressing the first two on the covariate ranks by least squares, and correlating the residuals. Reported *P* values use degrees of freedom reduced by the number of covariates and assume independent observations. Because measurements cluster within ovaries, this assumption is not met, so the *P* values are used for ranking rather than as calibrated error rates.

Ovary identity enters as indicator variables. Ovaries are nested within age group, so ovary and age cannot both be included in the pooled model; the pooled model uses age, and the within-ovary model uses ovary identity. Within each age stratum, ovary identity is estimable and is included. Five statistics are reported per gene: pooled with depth and age held constant, pooled with depth and ovary held constant, within young, within aged, and the difference between the two age strata.

Correlations were computed only on measurements in which the gene had at least one raw count. At a median depth of 21,572 counts per measurement this is a far weaker conditioning than the bin-level case in S9, though the operation is the same.

The interaction test compares the two within-stratum correlations by Fisher *z* transformation with the variance of the transformed Spearman correlation approximated as 1.06/(*n* − 3). At the sample sizes here any difference exceeding the stated effect-size threshold yields a nominal *P* value of order 10⁻⁷, so the reported count is determined by the effect-size threshold and not by the significance criterion.

Genes were tested if detected in at least 10% of measurements with at least 20 total counts, applied within the scope of each compartment analysis; 11,255 of 19,053 genes passed at whole-tissue level. Associations were retained as descriptive candidates when Benjamini-Hochberg-adjusted *P* values were < 0.05 and |ρ| ≥ 0.1. Grid points are spatially clustered within ROIs and ovaries, so these adjusted P values do not provide calibrated animal-level false-discovery control; robustness was assessed by leave-one-ovary-out analysis. Effect sizes are interpreted against the composition ceiling reported in S11.

Every correlation was recomputed with each ovary removed in turn, and the range and smallest absolute value across folds are reported. Ranked mechanical variance is partitioned into between-ovary and within-ovary components for each parameter, since the within-ovary test uses only the latter.

#### S11. Registration positive controls

Four analyses assessed internal reconstruction, spatial continuity, and coarse anatomical concordance between the registered datasets. These analyses do not quantify pointwise registration error or exclude local misregistration. The first two do not use the transcriptome.

##### Spatial continuity of the mechanical map

Each measurement’s log-transformed modulus was correlated against the mean of its four nearest neighbors within the same slice. Pooled ρ was +0.753, and +0.396 to +0.672 across the eight ovaries individually, at a median nearest-neighbor separation of 30.3 to 51.1 µm per slice.

##### Coarse anatomical concordance in stiffness after registration

Across all measurements, modulus differed by compartment (H = 85.2, P = 2.38 × 10⁻¹⁸). Compartment representation is strongly age-dependent, with corpus luteum making up 59% of young measurements and 6% of aged, so this pooled test partly reflects a difference between age groups. Rank order differed between age groups, consistent with the age-stratified result reported in the Online Methods, in which follicles are the stiffest compartment in young tissue and no compartment differs within aged tissue.

##### Association of compartment composition with stiffness

For each measurement, the fraction of its bins assigned to each compartment and each cell cluster was correlated with the mechanical parameter, with ovary identity held constant. The strongest association across all 38 annotated populations and the four compartments reached ρ = +0.056 (P = 0.003, FDR = 0.045), and three features passed correction. This represents the largest univariable composition-stiffness association observed under this analysis, rather than a general upper bound on compositional contributions.

##### Ovary-level exploratory association

Median stiffness per ovary was related to ovary-level pseudobulk expression at *n* = 8, across 14,818 genes. No gene could reach the corrected threshold at this sample size, so the result is reported as a scale rather than as a test.

Controls A and B both passed establishing that the mechanical map is spatially structured and that stiffness tracks transcriptomically defined architecture. Control C is a ceiling rather than a registration test: it bounds how much of the local stiffness variation cell composition can account for, and that ceiling is low. Control D is reported as a scale rather than a test, since with eight ovaries the minimum exact two-sided Spearman *P* value without ties is 4.96 x 10^-5^ which remains above the first Benjamini-Hochberg threshold of 3.4 x 10^-6^ for 14,818 tests. Control code is provided as Supplementary Code 16.

#### S12. Software and code

Analysis code is provided as Supplementary Code 1 to 18 (usage summarized in manifest README.txt), comprising nanoindentation force-curve processing and quality control (Supplementary Code 1 to 4), stiffness statistics (Supplementary Code 5), stiffness map generation (Supplementary Code 6), spatial transcriptomic bin size comparison (Supplementary Code 7), multi-sample integration and cell identity annotation (Supplementary Code 8), assignment of 4 µm transcriptomic bins to 50 µm nanoindentation grid points (Supplementary Code 9), pseudobulk differential expression and gene set enrichment (Supplementary Code 10), curation of gene set enrichment plots (Supplementary Code 11), stiffness-expression correlation analysis (Supplementary Code 12), merging of correlation results (Supplementary Code 13), spatial plotting (Supplementary Code 14), the sequencing depth diagnostic (Supplementary Code 15), registration positive controls (Supplementary Code 16), tilt correction of post-nanoindentation confocal stacks in Fiji (Supplementary Code 18), and an extras folder (Supplementary Code 17) containing elastic net regression of stiffness on cell type composition, which was run on this dataset, explained almost none of the variation, and is provided for users to test on other tissue or disease models,. Random seeds are fixed throughout (RANDOM_STATE 42). Parallel and serial execution paths were verified to produce identical output. Versions: Python 3.13.13 with NumPy 2.4.6, Pandas 2.3.3, SciPy 1.17.1, scikit-learn 1.9.0, statsmodels 0.14.6, patsy 1.0.2, Matplotlib 3.11.0, seaborn 0.13.2 and gseapy 1.3.0; R v4.5.1 with Seurat v5.3.1, BPCells v0.3.1, SeuratWrappers 0.4.0, Banksy 1.6.0, harmony 1.2.4, SpatialExperiment 1.20.0, SummarizedExperiment 1.38.1, scran 1.38.1, scater 1.38.1, clustree 0.5.1, Matrix 1.7.4, future 1.68.0, arrow 22.0.0, data. table 1.17.8, DESeq2 1.50.2, apeglm 1.32.0, fgsea 1.36.2 and msigdbr 26.1.0. Image processing used Fiji (ImageJ 1.53q, Java 1.8.0_322) and Imaris 10.2.0 (Oxford Instruments). Image alignment and ROI annotation were performed in SpatialX v2026-08-03 (BioTuring Inc).

## Extended Data Tables

\**Tables to be uploaded upon journal submission*

**Extended Data Table 1.** VisiumHD sequencing and capture metrics. Space Ranger summary for each of the four capture areas, covering read depth, barcode and UMI validity, Q30 base fractions, probe set mapping rates, and the number of squares and mean reads, genes and UMIs per square under tissue at 2, 8 and 16 µm.

**Extended Data Table 2.** Marker genes for cell identity annotation. One-versus-all Wilcoxon rank-sum results for each of the 38 annotated populations, ranked by a composite score combining effect size, detection specificity and statistical confidence, with average expression per cluster.

**Extended Data Table 3.** Differential expression between aged and young ovaries. Pseudobulk DESeq2 results computed per animal, with counts from both ovaries summed where an animal contributed a pair, across 16 ovaries from 10 animals (5 per age group), with one tab for whole tissue and one for each of the four compartments. Negative fold changes denote higher expression in young tissue. Both shrunken and unshrunken fold-change estimates are given.

**Extended Data Table 4.** Gene set enrichment between aged and young ovaries. Complete GSEA results on the ranked Wald statistic against the mouse-native MSigDB Hallmark, GO Biological Process, GO Cellular Component, GO Molecular Function and Reactome collections, with one tab for whole tissue and one for each compartment. Positive NES indicates enrichment in aged tissue.

**Extended Data Table 5.** Ovarian Mechanomics gene set. All 1,854 curated symbols with the route by which each entered the list, matrisome membership and category, ligand-receptor role, and full partner lists.

**Extended Data Table 6.** Ovarian Mechanomics summary. The same 1,854 symbols with age-associated differential expression across whole tissue and each compartment, and partial Spearman correlations with stiffness. Of these, 1,724 are present in the expression matrix and 1,039 passed the detection filter for correlation in at least one region. A blank cell indicates a gene absent from the matrix or not tested in that analysis, as given by the in_expression_matrix and tested_correlation columns.

**Extended Data Table 7. Sample, acquisition and region of interest identifiers.** One row per nanoindentation region of interest, with rows for ovaries and serial sections profiled by spatial transcriptomics alone. Study ID identifies the animal and is the label used in the figures. Contralateral indicates whether the paired ovary from the same animal was also profiled and whether it was indented. ST section gives the serial cryosection transferred to the Visium HD capture area. Storage is the interval between fresh freezing and cryosectioning. Indentations passing QC is the number of measurements within that region of interest retained after nanoindentation quality control, and grid points analyzed is the subset matched to at least five transcriptome bins. Sixteen ovaries from ten animals were profiled by spatial transcriptomics, of which eight ovaries from six animals carry matched nanoindentation on the adjacent surface, giving 21 regions of interest and 2,771 analyzed grid points.

## Notes

### Competing Interest Statement

The authors have declared no competing interest.

### Summary of Updates

Since posting the original version, we refined the paired-surface registration and alignment quality criteria. These revisions changed a number of retained stiffness-expression measurements and some downstream associations; all values and analyses in this manuscript reflect the revised computational workflow and supersede those reported in the initial preprint.

